# Nuclear cytoglobin associates with HMGB2 and regulates DNA damage and genome-wide transcriptional output in the vasculature

**DOI:** 10.1101/2023.05.10.540045

**Authors:** Clinton Mathai, Frances Jourd’heuil, Le Gia Cat Pham, Kurrim Gilliard, Dennis Howard, Joseph Balnis, Ariel Jaitovich, Sridar V. Chittur, Mark Rilley, Ruben Peredo-Wende, Ibrahim Ammoura, Sandra J. Shin, Margarida Barroso, Jonathan Barra, Evgenia Shishkova, Joshua J. Coon, Reynold I. Lopez-Soler, David Jourd’heuil

## Abstract

Identifying novel regulators of vascular smooth muscle cell function is necessary to further understand cardiovascular diseases. We previously identified cytoglobin, a hemoglobin homolog, with myogenic and cytoprotective roles in the vasculature. The specific mechanism of action of cytoglobin is unclear but does not seem to be related to oxygen transport or storage like hemoglobin. Herein, transcriptomic profiling of injured carotid arteries in cytoglobin global knockout mice revealed that cytoglobin deletion accelerated the loss of contractile genes and increased DNA damage. Overall, we show that cytoglobin is actively translocated into the nucleus of vascular smooth muscle cells through a redox signal driven by NOX4. We demonstrate that nuclear cytoglobin heterodimerizes with the non-histone chromatin structural protein HMGB2. Our results are consistent with a previously unknown function by which a non-erythrocytic hemoglobin inhibits DNA damage and regulates gene programs in the vasculature by modulating the genome-wide binding of HMGB2.

## Introduction

Hemoglobins are essential components of the cardiovascular system where they transport and store molecular oxygen and regulate nitric oxide (NO) and nitrite (NO_2-_) signaling^1^. In addition to myoglobin and hemoglobin-α, the hemoglobin paralog cytoglobin is expressed in medial vascular smooth muscle (VSM) cells, in human and rodent vessels^2–4^. Despite detailed characterization of the structure and biochemical properties of cytoglobin, its function and mechanism of action are still unclear. The role of cytoglobin in oxygen transport and storage in vessels is doubtful because the intracellular concentration of cytoglobin is too low to affect oxygen diffusion rates. In contrast, the NO dioxygenase activity of cytoglobin that inactivates NO to nitrate (NO_3-_) is relatively fast and facilitated by efficient reduction of cytoglobin in cells^4–7^. As such, cytoglobin has been proposed to regulate NO dependent-vasorelaxation and NO-dependent cytotoxicity in vessels during conditions associated with increased inducible NO synthase (NOS2) expression^4, 8, 9^. More recently, we found that isolated vessels from cytoglobin knockout mice also displayed decreased rates of hydrogen peroxide consumption, suggesting a role for cytoglobin as a peroxidase in the vasculature^10^. In models of vascular injury, we proposed that loss of cytoglobin limited the survival capacity of a cell population that expresses cytoglobin 2-3 days after initiation of the injury^9^. The loss of cytoglobin coincided with an increase in caspase 3 activity upon injury of the rat carotid artery, suggestive of increased apoptosis^9^. Similarly, loss of cytoglobin *in vitro* sensitized human vascular smooth muscle to staurosporine-mediated cell death^9^. In another model that combined the mouse carotid artery ligation model with global deletion of cytoglobin, we still observed an inhibition in neointima formation but in this case, no evidence for cell death was found^9^. While it is tempting to attribute these effects to specific antioxidant properties derived from cytoglobin expression, direct evidence is still lacking. Particularly interesting to us were past reports indicating broad gene regulation following manipulation of cytoglobin levels^11, 12^, suggesting that alternative mechanisms might be at play. In addition, loss of cytoglobin *in vivo* or *in vitro* has been associated with an increase in DNA damage^13–15^. This raised the possibility that cytoglobin could regulate gene expression and DNA damage in vessels and this could contribute to the observed phenotypic changes associated with loss of cytoglobin after vascular injury. In the present study, we combined transcriptional and protein profiling with functional validation *in vitro* and *in vivo* to reveal an unexpected mechanism by which cytoglobin regulates gene expression and DNA repair pathways. We delineate a model where NOX4 and hydrogen peroxide promote the nuclear translocation of cytoglobin. In the nucleus, cytoglobin heterodimerizes with the non-histone DNA interacting protein HMGB2 and serves as a regulator of HMGB2. To our knowledge, this is the first demonstration of a mammalian globin regulating transcriptional output and cell fate.

## Results

### Deletion of cytoglobin accelerates the downregulation of smooth muscle contractile genes and increases DNA damage in injured vessels in mice

An abnormal hyperplastic response characterized by smooth muscle cell de-differentiation to proliferative and inflammatory phenotypes occurs in atherosclerotic and stenotic vessels^16^. Past studies from this laboratory showed that 4 weeks after complete ligation of the left common carotid artery, cytoglobin knockout mice displayed no evidence of neointimal hyperplasia in contrast to the wildtype^9^. Although we showed that the decrease in neointima formation following cytoglobin silencing in the rat carotid artery angioplasty model was associated with increased smooth muscle apoptosis^9^, the underlying reasons for the absence of neointima in the mouse ligation model was not investigated. To gain further insights into the mechanism by which cytoglobin regulates vascular injury, cytoglobin knockout mice (Cygb KO, **Fig. 1a**) and their wild-type (Cygb WT) littermates were subjected to carotid artery ligation, the uninjured right and injured left common carotid arteries were collected for bulk transcriptomics analysis. Sampling 3-day post-injury was used for these studies because early transcriptional changes of VSM dedifferentiation are most evident at this time^17^. We first confirmed loss of cytoglobin protein following deletion and 3 days post-ligation (**Fig. 1a**) and cellular enumeration indicated no difference in medial and adventitial cell numbers in the wild-type and the knockout littermates (**Supplementary Fig. 1**). Differential gene expression analysis to isolate changes that occur exclusively in the injured vessels of cytoglobin knockout mice reveals a total of 2993 genes uniquely changed with 1086 genes upregulated and 1907 genes downregulated (**Fig. 1b and c**). Among primary downregulated gene sets, gene ontology pathway analysis identified processes associated with differentiated smooth muscle cells (contractile fiber, muscle structure development, and muscle contraction) and extracellular matrix production (**Fig. 1c**). Transcriptionally downregulated smooth muscle contractile genes included phosphodiesterase 5a (Pde5a), myosin light chain kinase (Mylk), smoothelin (Smtln), leiomodin 1 (Lmod1), myosin heavy chain 11 (Myh11), calponin (Cnn1), and the serum responsive factor (Srf) co-regulator myocardin (Myocd; **Fig. 1d**). Consistent with the RNA Seq results, we found a statistically significant decrease in immunostaining for ACTA2 in the injured vessel from cytoglobin knockout mice compared to wild type (**Fig. 1e)**. Our results indicate that cytoglobin plays a role in regulating the expression of smooth muscle differentiation markers *in vivo* in response to injury.

**Figure 1.**
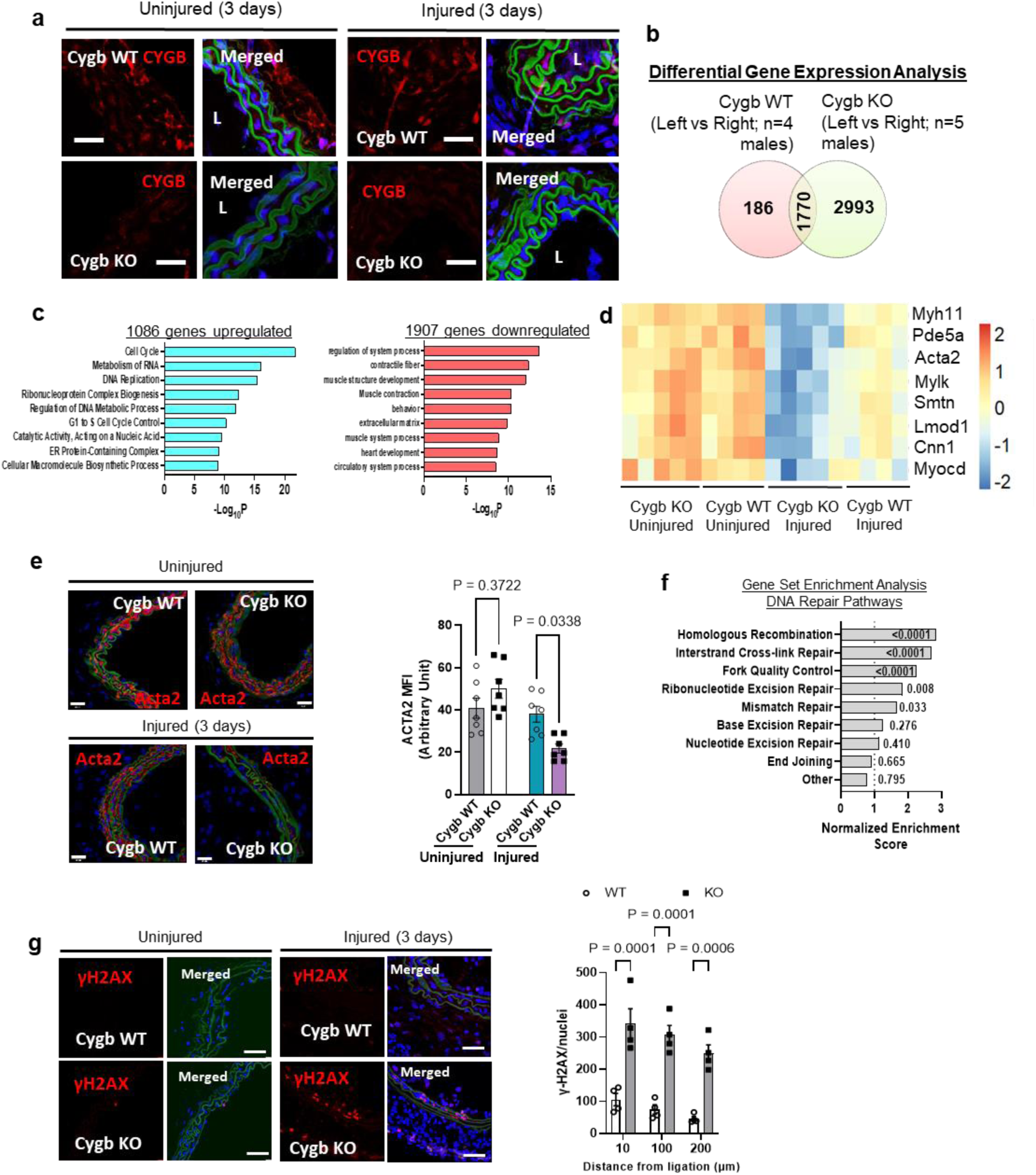
Global deletion of cytoglobin in the mouse accelerates the decrease in smooth muscle contractile genes and activation of DNA repair pathways in the carotid artery ligation model. (**a**) Representative immunofluorescence staining of the right (uninjured) and left (injured) common carotid arteries of cytoglobin wildtype (WT) and knockout (KO) littermate mice 3-days post-ligation, harvested and analyzed for cytoglobin content (red – CYGB, blue – DAPI, green – autofluorescence marking elastic lamina). Cytoglobin immunostaining was evident in the media and adventitia (top) and lost in the KO mice; scale bar, 40 µm. (**b**) Results from bulk RNA-Seq analysis of the entire left (injured) and right (uninjured) common carotid arteries from mice that underwent left common artery ligation for 3 days. 2993 genes differentially changed in the injured cytoglobin knockout mice were isolated from differential gene expression analysis of wildtype injured (Left) and uninjured (Right) vessels (n=4) and cytoglobin knockout injured and uninjured (n=5; 1.5 – fold change FDR < 0.05). **(c)** Top down-and up-regulated pathways from gene ontology analysis of the 2993 genes altered in the differential gene expression analysis. **(d)** Heat map comparing smooth muscle cell contractile genes in CYGB-WT and CYGB-KO mice 3 days after left common carotid artery ligation (color scheme represents the z-score across each individual gene. **(e)** Representative immunofluorescence images and fluorescence quantitation from CYGB-WT and CYGB-KO carotids probing for ACTA2, 3 days after ligation; scale bar, 50 µm. **(f)** Gene set enrichment analysis based on a curated set of 197 DNA damage factors associated with DNA repair pathways; The number for each bar represents the nominal P value for each pathway. **(g)** Indirect immunofluorescence staining for the marker of DNA repair activation γH2AX (phosphorylated histone 2AX) of uninjured (right common) and injured (left common) mouse carotid arteries 3-days post ligation (red – γH2AX, blue – DAPI, green – autofluorescence marking elastic lamina); scale bar, 100 µm. Right panel, quantitation of results. For statistical analysis throughout the figure, results were analyzed by two-way ANOVA, followed by Tukey’s post hoc test.

An important phenotypic attribute previously associated with the loss of cytoglobin *in vivo* is an increase in DNA damage^14^. To determine whether loss of cytoglobin in the mouse model of carotid ligation was also associated with changes consistent with increased DNA damage, we first used a curated set of 197 DNA damage repair and signaling factors grouped according to their known association with specific DNA repair pathways^18^. We examined which groups might be enriched in the cytoglobin knockout mice compared to their wild type littermates 3-days post ligation in the carotid arteries by gene set enrichment analysis (**Fig. 1f**). We found that genes preferentially associated with “homologous recombination”, “inter-strand crosslink repair”, “fork quality control” and “mismatch repair” were statistically significantly increased. Next, we performed some immunofluorescence staining for the phosphorylated form of histone H2AX called γ-H2AX, a common approach to assess DNA damage^19^. We found no evidence of γ-H2AX staining in uninjured vessels in cytoglobin knockout mice over background levels obtained in the wild type mice (**Fig. 1g**). However, injured vessels in the knockout mice had a statistically significant increase in γH2AX staining compared to wild type littermates (**Fig. 1g**). Taken together, these results indicate that cytoglobin is required for DNA damage repair during conditions associated with injury.

### Cytoglobin regulates gene expression and DNA repair pathways in human vascular smooth muscle cells in vitro

To investigate the functional significance of cytoglobin in human vascular smooth muscle cells (SMC), we used siRNA mediated knockdown of cytoglobin and performed a differential gene expression analysis in subcultured human aortic vascular smooth muscle cells. Electroporation of siRNAs targeting cytoglobin decreased cytoglobin mRNA transcripts by approximately 70% and protein expression by approximately 60% with no change in the scrambled controls (**Fig. 2a and 2b**). Differential gene expression analysis revealed broad changes in gene expression following cytoglobin silencing (**Fig. 2c**). Post-analysis of the differentially expressed gene set was performed using Ingenuity Pathway Analysis (QIAGEN Inc.). The top scoring networks were related to cell cycle, cellular assembly, DNA replication, recombination, and repair. Further analysis probing for diseases and functions yielded relationships to cardiovascular system and function as well as cell death and survival. (**Fig. 2d and e**). The IPA analysis also identified DNA repair as a key feature associated with cytoglobin silencing in these cells. To confirm the occurrence of DNA damage, we immuno-stained human vascular smooth muscle cells for γ-H2AX. We show that 100 µM and 200 µM treatments of hydrogen peroxide for ten minutes were sufficient to induce γ-H2AX formation (**Supplementary Fig. 2**). The siRNA mediated knockdown of cytoglobin significantly increased γH2AX formation in comparison to cells treated with a non-targeting siRNA (**Fig. 2f**). These results were further validated by COMET assay and comet tail moment was significantly increased in cells silenced for cytoglobin (**Fig. 2g**). Altogether, these results show that cytoglobin is involved in a cell autonomous fashion in DNA repair pathways and its expression inhibits hydrogen peroxide induced DNA damage.

**Figure 2.**
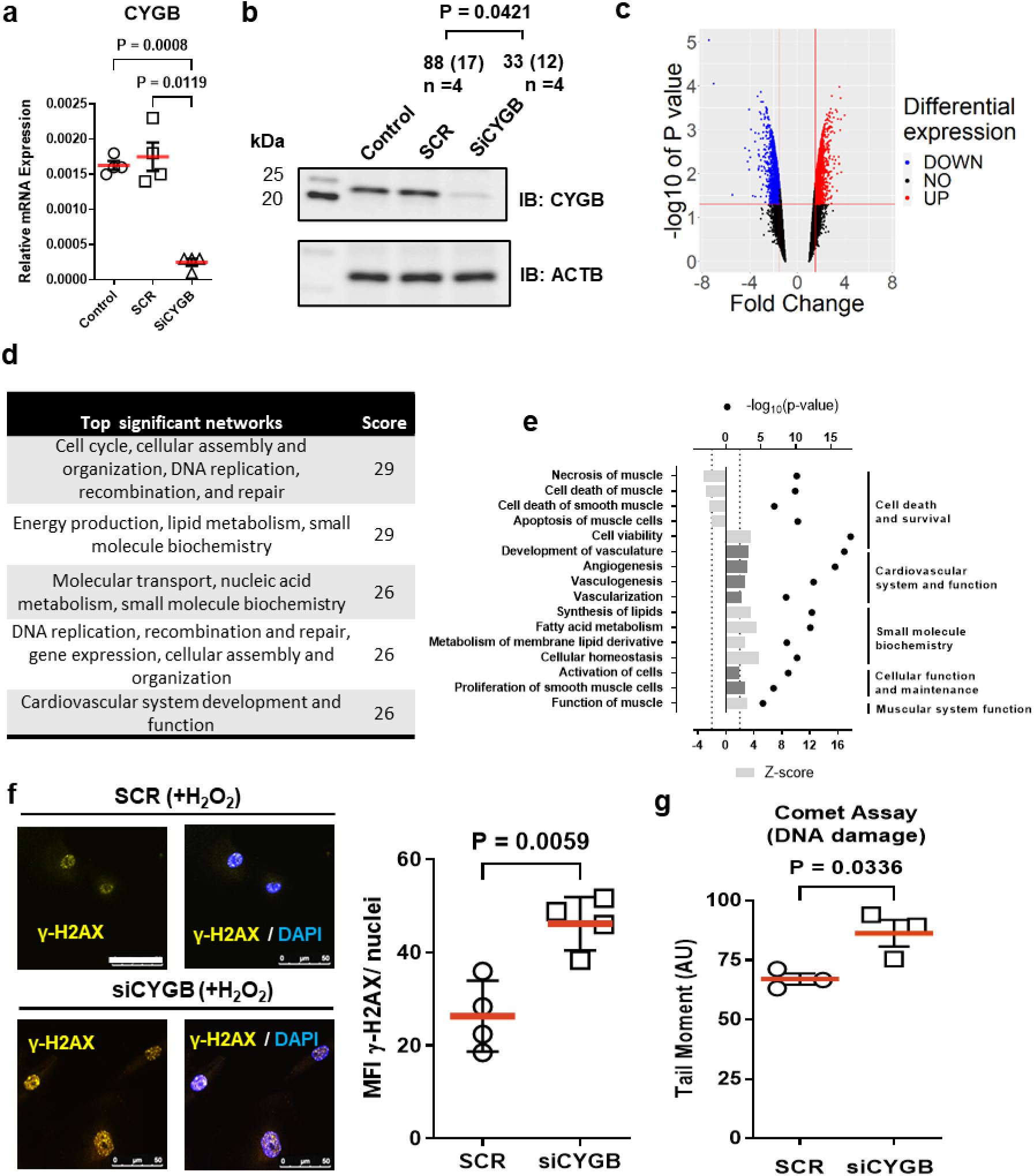
Decrease in cytoglobin expression in human smooth muscle cells alters gene expression and increases hydrogen peroxide-induced DNA damage *in vitro*. (**a**) Quantitative reverse transcriptase polymerase chain reaction (qRT-PCR) results showing the relative expression of cytoglobin (CYGB) in sub-cultured human vascular smooth muscle cells following silencing of cytoglobin. Human smooth muscle cells were electroporated with scrambled SiRNA (SCR) or siRNA mix targeting CYGB (SiCYGB). Results from non-transfected cells are also shown (Control). **(b)** Western blot shows protein levels of CYGB using conditions described in (a); values represent the mean and (SEM) of the percentage change in CYGB immunoreactivity compared with non-transfected conditions (Control) using ACTB as an internal reference (n=4). **(c)** Volcano plot of the differential gene expression analysis following silencing of CYGB in human vascular smooth muscle cells. **(d)** Top 5 significant networks for disease and function from transcriptomic results following CYGB silencing in sub-cultured human vascular smooth muscle cells was generated with QIAGEN Ingenuity Pathway Analysis (IPA). **(e)** The IPA regulation z-score was used to identify diseases and functions that are either increased (z-score>= 2) or decreased (z-score <=-2) and the p-value calculated with the Fischer’s exact test indicates the likelihood that the association between a set of genes and a biological function is significant. **(f)** Representative immunofluorescence images of yH2AX (yellow) without (left) or with (right) DAPI staining in human vascular smooth muscle cells electroporated with scrambled SiRNA (SCR) or siRNA mix targeting CYGB (SiCYGB) for 72 hours followed by treatment with hydrogen peroxide for 10 min; scale bar 50 µm. The right-side figure is quantification of results from 4 independent experiments. **(g)** Comet assay was performed under alkaline unwinding/neutral electrophoresis conditions and quantified (n = 3). Throughout the figure, data are presented as mean +/-SEM; p values were determined by Student’s t test or one way ANOVA. Red bars represent the mean for each condition.

### Cytoglobin is trafficked to the nucleus through NOX4 and hydrogen peroxide

The cytoplasmic and nuclear association of cytoglobin has been recognized for many years^20^. Its functional significance and underpinning mechanisms are, however, unknown. In the present work, confocal images revealed temporal changes in cytoglobin localization in human vascular smooth muscle cells when changing from low to high serum-containing media. (**Fig. 3a**). Kinetic analysis revealed that nuclear cytoglobin increased within minutes of addition of growth media, saturated within ∼ 7 hours, and was sustained for at least 24 hours (**Fig. 3b**). This was reversed following replacement of the growth media with starvation media and maximal nuclear depletion was achieved within 5 hours (**Supplementary Fig. 3**). To determine whether the nuclear accumulation of cytoglobin was a regulated or passive process, growth media stimulated cells were treated with ivermectin, an importin-α/β inhibitor^21^. This decreased the nuclear to cytoplasmic ratio for cytoglobin content, consistent with importin-dependent nuclear import of cytoglobin during growth media stimulation (**Fig. 3c**). In contrast, pharmacological inhibition of Chromosomal Maintenance 1 (CRM1) – one of the main nuclear exportins – by leptomycin B^22^ caused an increase in the ratio of nuclear to cytoplasmic cytoglobin in serum starved cells (**Fig. 3c**). Overall, these results are consistent with a model where cytoglobin exclusion from the nucleus in quiescent smooth muscle cells is maintained through rapid CRM1-dependent nuclear export. Conversely, the nuclear import of cytoglobin in growth media stimulated cells appears to be dependent on importin-α/β1 mediated trafficking.

**Figure 3.**
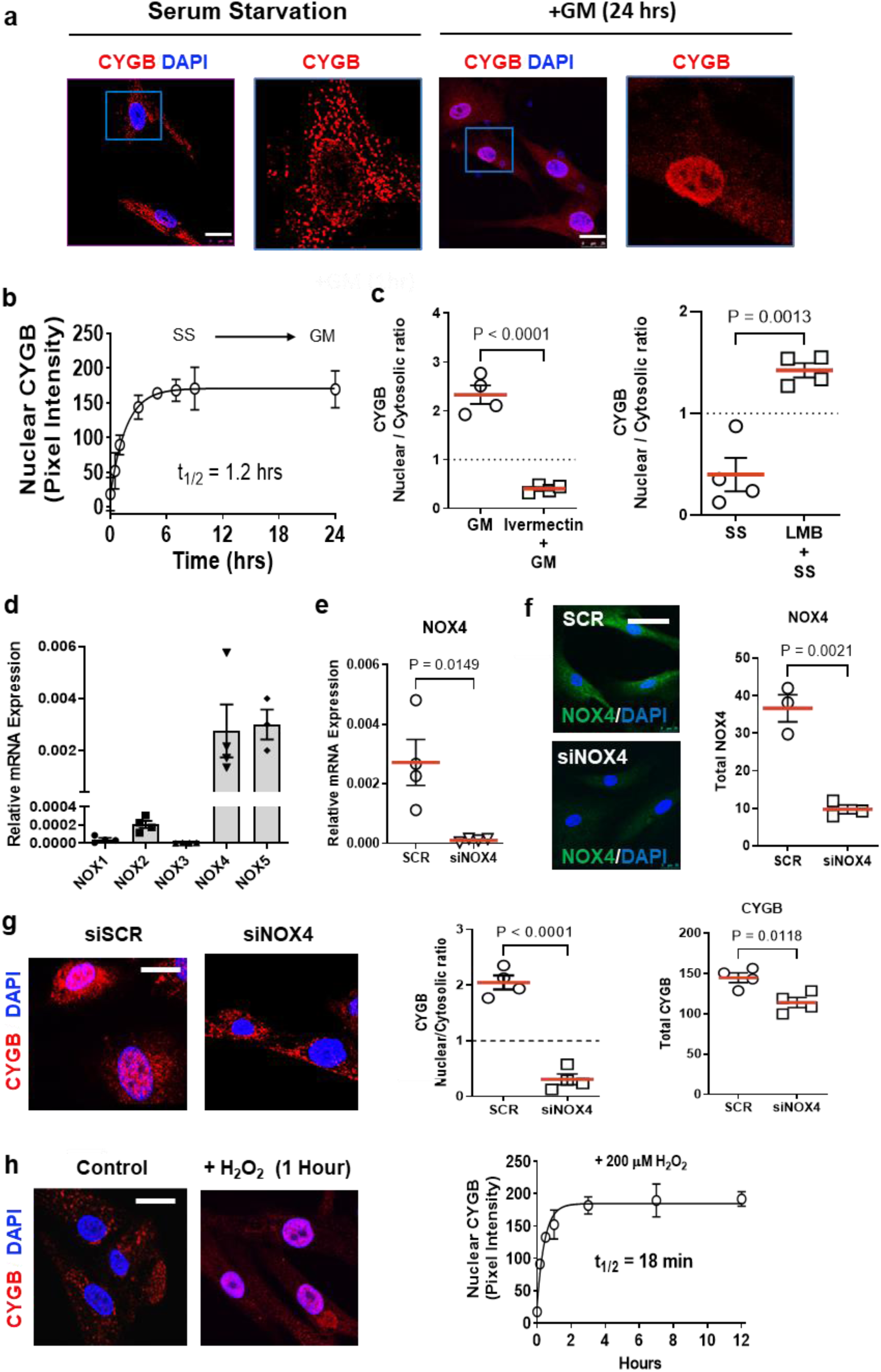
The hydrogen peroxide producing NADPH oxidase 4 (NOX4) promotes the active trafficking of cytoglobin to the nucleus in human vascular smooth muscle cells. **(a)** Representative confocal immunofluorescence images of sub-cultured human vascular smooth muscle cells following 0-, and 24-hours stimulation with 5% serum containing growth media (Blue – DAPI, red – CYGB). Cells were then fixed and analyzed by indirect immunofluorescence using an antibody against cytoglobin; scale bars, 25 µm. **(b)** Time course of cytoglobin nuclear import following the addition of 5% serum containing growth media. The solid line represents the non-linear regression obtained from a single exponential model yielding a half-life 1.2 hours; n = 4. **(c)** Quantitative analysis of the ratio of nuclear to cytosolic cytoglobin (CYGB) pixel intensity after pre-treatment with ivermectin (nuclear import inhibitor, left panel) or Leptomycin B (right panel, LMB – nuclear export inhibitor). A ratio greater than 1 is indicative of nuclear accumulation. **(d)** Quantitative reverse transcriptase polymerase chain reaction (qRT-PCR) results showing the relative expression for NADPH oxidase (NOX) isoforms in sub-cultured human vascular smooth muscle cells. **(e)** Quantitative reverse transcriptase polymerase chain reaction (qRT-PCR) results showing the relative expression of NADPH oxidase 4 (NOX4). Human smooth muscle cells were electroporated with scrambled SiRNA (SCR) or siRNA mix targeting NOX4. Each individual symbol represents an independent experiment. **(f)** Left panel, representative indirect immunofluorescence pictures of human vascular smooth muscle cells stained for NOX4 following silencing of NOX4 as described in panel (e); bar scale, 50 µm. Right panel, quantitation of experiments shown in left panel. **(g)** Left panel, representative indirect immunofluorescence pictures of human vascular smooth muscle cells stained for cytoglobin following silencing of NOX4 as described in panel (e); bar scale 20 µm. Center panel, quantitative pixel analysis of the ratio of nuclear to cytosolic cytoglobin (CYGB) pixel intensity following silencing of NOX4. Right panel, quantitative pixel analysis of total cytoglobin (CYGB) pixel intensity following silencing of NOX4. **(h)** Right panel, representative indirect immunofluorescence pictures of serum-starved (control) human vascular smooth muscle cells stained for cytoglobin following treatment with 200 µM hydrogen peroxide (+H2O2) for 1 hour. Right panel, time course of cytoglobin nuclear import following the addition 200 µM hydrogen peroxide as described for left panel. The line represents the non-linear regression derived from a single exponential model yielding a half-life 18 min; n = 4; p values were determined by Student’s t test or one way ANOVA, followed by Tukey’s post hoc test. Red bars represent the mean for each condition and error bars represent SEM.

Next, we tested the hypothesis that cytoglobin nuclear translocation depends on a redox signal. Pretreatment of growth media stimulated cells with the antioxidant N-acetylcysteine (NAC) inhibited the nuclear translocation of cytoglobin (**Supplementary Fig. 4**). Similar results were obtained using the flavoenzyme inhibitor diphenyleneiodonium (DPI; **Supplementary Fig. 4**). However, in this case total cytoglobin content was also decreased. Analysis of the relative content of NADPH oxidase (NOX) transcripts by QPCR in human aortic vascular smooth muscle cells suggested that the primary isoforms expressed in these cells were NOX4 and NOX5 (**Fig. 3d**). We decreased cellular levels of these two proteins individually through electroporation of gene-specific siRNAs and verified specific silencing through QPCR and immunofluorescence (**Fig. 3e, f and Supplementary Fig. 5**). We found that silencing NOX4 and not NOX5 inhibited cytoglobin nuclear accumulation in response to growth media (**Fig. 3g and Supplementary Fig. 5**). Treatment of serum-starved cells with hydrogen peroxide was sufficient to induce the nuclear translocation of cytoglobin (**Fig. 3h**) with an EC_50_ of 87 µM (**Supplementary Fig. 6**). Overall, these results are consistent with the accumulation of cytoglobin in the nucleus in response to growth factor-mediated increase in hydrogen peroxide production through NOX4.

### Expression of nuclear cytoglobin is sufficient to inhibit DNA damage from hydrogen peroxide, but its hydrogen peroxide scavenging activity is dispensable

The aim of the next experiments was to define the mechanism by which cytoglobin regulates DNA repair pathways. In the first set of experiments, we wanted to determine whether cytoglobin nuclear association was required to inhibit hydrogen peroxide induced DNA damage. Terminal ends of proteins may harbor sequences regulating nuclear import and export. The primary structure of cytoglobin contains two unique N-and C-terminal ends (**Fig. 4a**), and we hypothesized that these may provide regulation over the subcellular localization of cytoglobin. We used N-and C-terminal end deletion mutants (ΔN or ΔC) and ectopically expressed them in HEK293 cells (**Fig. 4a**). We did not use the ΔNΔC double mutant because levels of cytoglobin expressions were inconsistent across experiments (**Fig. 4a**). Indirect immunofluorescence analysis showed exclusion of the ΔC deletion mutant from the nucleus, while the ΔN deletion mutant localized in the nucleus (**Fig. 4b and c**). This contrasted with the more diffuse nuclear and cytoplasmic association of full length cytoglobin. We also treated these clones with hydrogen peroxide, assessed subcellular localization of cytoglobin, and stained for γ-H2AX. There was no additive effect related to nuclear accumulation and hydrogen peroxide treatment, except for the ΔN variant, which was already nuclear (**Fig.4c**). Additionally, we saw statistically significant decreases in γH2AX staining in the hCYGB and ΔN clones compared to the empty vector control (**Fig. 4d and 4e**), both of which can localize into the nucleus. The ΔC mutant, which exclusively associates with the cytosol, had significantly higher γ-H2AX staining (**Fig. 4d and 4e**). To verify that the ΔC mutant variant was still functional, we fused a canonical nuclear localization sequence (NLS) to our ΔC clone (ΔC-NLS). These cells were then treated with hydrogen peroxide to induce DNA damage. In contrast to the cytosolic ΔC variant, we found complete inhibition of DNA damage based on the significant decrease in γ-H2AX staining (**Fig. 4f and 4g**). In agreement, there was a significant decrease in COMET tail moment in ΔC-NLS cells compared to ΔC cells (**Fig. 4h**). Taken together, these results demonstrate that cytoglobin is required in the nucleus to inhibit DNA damage and affect DNA repair pathways.

**Figure 4.**
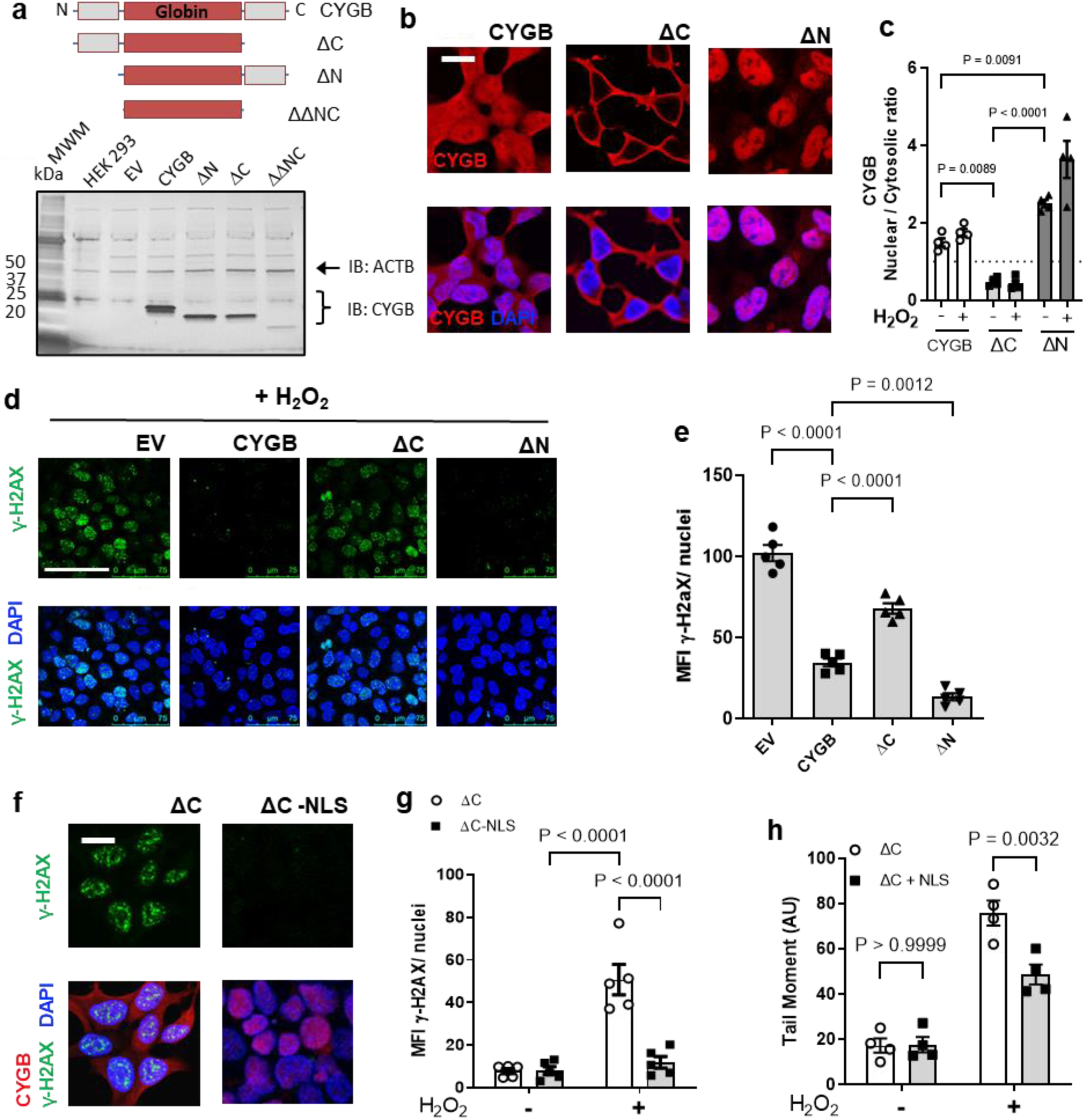
Nuclear cytoglobin is sufficient to inhibit DNA damage. (**a**) Top panel, schematic of the primary sequence of cytoglobin composed of a central globin domain and two unique 20 amino acid C-and N-terminal ends. Cytoglobin expression plasmids were generated for full length cytoglobin, and the N and C terminal truncated variants. Expression from the different plasmid in HEK293 was examined by Western blot. The double truncated variant did not express consistently. ACTAB is shown as a loading control. **(b)** Indirect immunofluorescence staining using an antibody directed against cytoglobin (red). **(c)** Quantitative analysis of the ratio of nuclear to cytosolic cytoglobin (CYGB) pixel intensity in HEK293 with plasmids expressing cytoglobin or empty plasmid (pcDNA) and treated with hydrogen peroxide. **(d)** Representative immunofluorescence images of γ-H2AX (green) staining in HEK293 expressing cytoglobin or empty plasmid following treatment with 200 µM hydrogen peroxide. **(e)** Quantitation of the mean fluorescence intensity of γ-H2AX per nuclei following 200 µM hydrogen peroxide treatment. **(f)** Representative immunofluorescence images of HEK293 cells expressing cytoglobin with C-terminal truncation and C-terminal truncation + nuclear localization sequence (NLS) exposed to 200 μM hydrogen peroxide and stained for γ-H2AX (green), DAPI (blue) and CYGB (red). **(g)** Quantitative analysis of γ-H2AX pixel intensity from results described in (f). Each point represents an independent experiment. **(h)** Quantitation of COMET tail moment from alkaline COMET assay in HEK C-terminal truncation and HEK C-terminal truncation + NLS exposed to 200 μM hydrogen peroxide. For statistical analysis throughout the figure, results were analyzed by one-way or two-way ANOVA followed by Tukey’s post hoc test for multiple comparisons.

### Cytoglobin affects DNA repair pathways and inhibits DNA damage independent of hydrogen peroxide scavenging

In a recent study, we found that cytoglobin can function as an antioxidant through scavenging of cytosolic hydrogen peroxide^10^. To test whether cytoglobin inhibits DNA damage through hydrogen peroxide scavenging in the nucleus, we expressed HyPer7 in HEK293 cells, a genetically encoded fluorescent probe specific for hydrogen peroxide detection^23^ (**Fig. 5a**). We used a HyPer7 construct tagged with a nuclear localization sequence (NLS) to measure hydrogen peroxide levels over time specifically in the nucleus. Cells expressing hCYGB showed a significant increase in the rate of hydrogen peroxide reduction in the nucleus. In contrast, we show that – despite inhibition of DNA damage (**Fig. 4g and h**) – the nuclear localized ΔC-NLS mutant showed no significant change in the rate of hydrogen peroxide degradation over time compared to the empty vector control (**Fig. 5b**). This forced us to conclude that the inhibitory effect of cytoglobin on DNA damage cannot be explained solely by its antioxidant activity. Several studies have shown that the majority of hydrogen peroxide handling occurs in the cytosol^23, 24^ and we suggest that the hydrogen peroxide scavenging activity of cytoglobin might be restricted to the cytosol. To confirm the inhibitory effect of cytoglobin on DNA damage independent of its antioxidant activity, we treated cells with 50 μM camptothecin for 1 hour and immunostained for γ-H2AX and CYGB. Camptothecin is a topoisomerase poison and non-oxidant inducer of DNA damage^25^. We found that nuclear cytoglobin (both ΔC-NLS and ΔN) decreased γ-H2AX staining and inhibit DNA damage (**Fig. 5c and d**). Overall, these results suggest that cytoglobin is required in the nucleus to affect gene programs related to DNA repair and mitigate both oxidant and replicative – induced DNA damage. Moreover, we show that these effects occur through a mechanism separate from the previously established oxidant scavenging activity of cytoglobin^10^.

**Figure 5.**
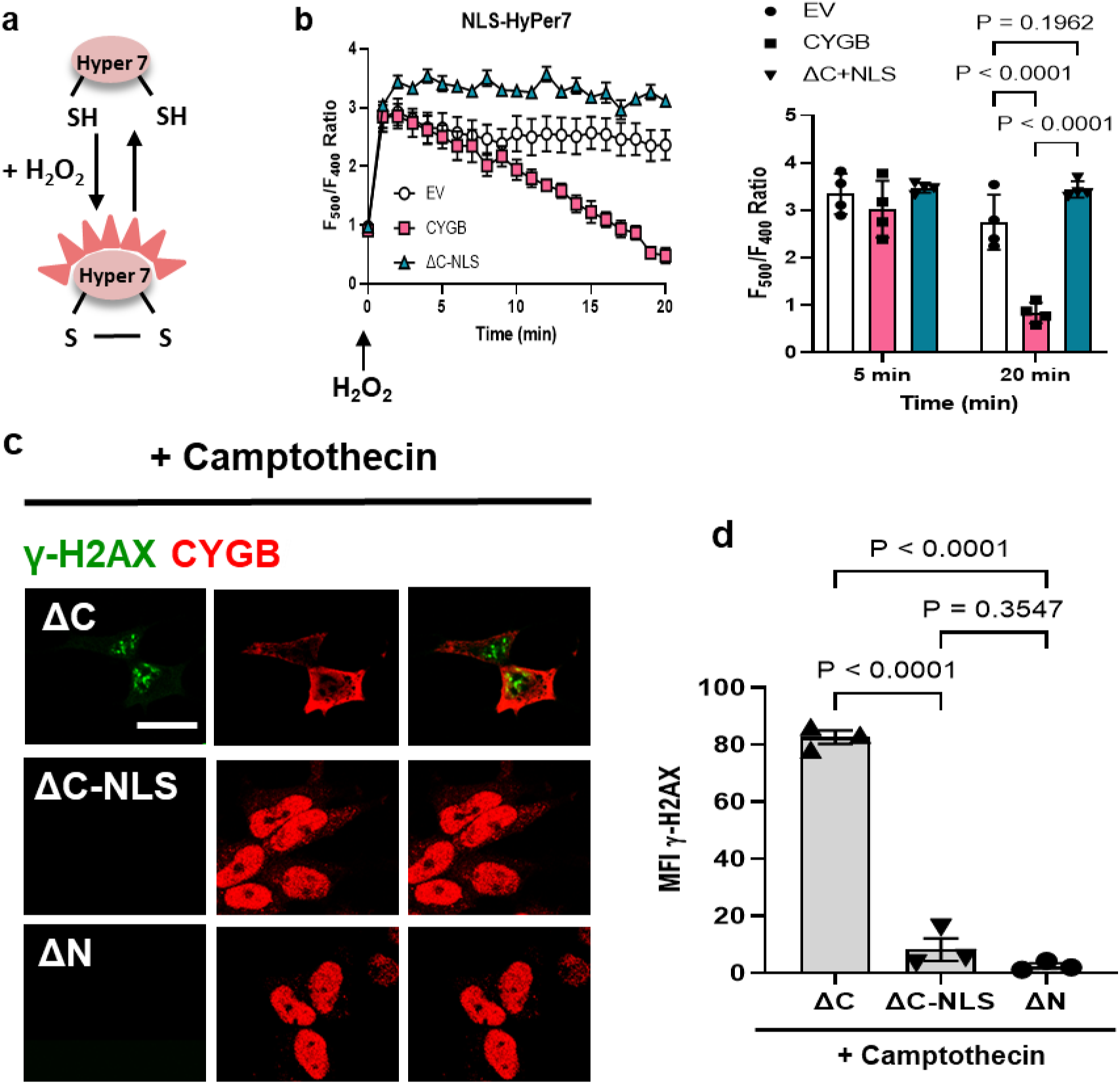
Cytoglobin inhibits DNA damage independent of its reductive activity. (**a**) Schematic representation of the hydrogen peroxide sensor Hyper7. It contains a pair of cysteine residues that can be reversibly oxidized by hydrogen peroxide leading to changes in its fluorescence properties **(b)** HEK293 cells stably transfected with pcDNA (empty vector), hCYGB (human cytoglobin) and ΔC-NLS (C-terminal truncation with nuclear localization sequence) plasmids were transiently transfected with HyPer7 with targeted NLS. Cells were then treated with 200µM bolus hydrogen peroxide and Hyper7 oxidation was measured over 20 minutes. Right panel, bar graph of fluorescence ratios for Hyper7, 5 and 20 min after the addition of hydrogen peroxide. **(c)** Representative immunofluorescence images of C-terminal truncated, C-terminal truncated +NLS and N–terminal truncated CYGB variants pre-treated with 50 μM camptothecin (CPT – topoisomerase inhibitor, oxidant-independent inducer of DNA damage) for 1 hour and immunostained for γ-H2AX (green) and CYGB (red); scale bar, 40µm. **(d)** Quantitative analysis of γ-H2AX pixel intensity. Each point represents one independent experiment. For statistical analysis throughout the figure, results were analyzed by two-way ANOVA, or one way ANOVA followed by Tukey’s post hoc test.

### Interactome analysis reveals heterodimerization of cytoglobin with nuclear proteins

To delineate mechanisms by which cytoglobin regulates DNA repair and gene expression, we analyzed the protein interactome of cytoglobin through immunoprecipitation coupled to mass spectrometry. To this end, pCMV6-tag (empty vector) or hCYGB-tag (wild type human cytoglobin) were expressed in HEK293 cells and lysates were obtained from both groups with and without treatment with 150 μM hydrogen peroxide for 10 min (**Fig. 6a**). Following immunoprecipitation and mass spectrometry, we identified a total of 87 protein interactors (at p<0.001, **Fig. 6b and Supplementary Tables 2 and 3**) among which, 31 were uniquely detected following hydrogen peroxide treatment. Of these 31 proteins, we curated a list of nuclear associated proteins and ordered them based on their fold-change following immunoprecipitation (**Fig. 6b**). This included Metastasis Associated 1 (MTA1), a structural component of the Nucleosome Remodeling and Deacetylase complex (NuRD)^26^, and High Mobility Group Box 2 (HMGB2), a non-histone chromatin structural protein^27^. We could not confirm the interaction between cytoglobin and MTA1, due to the lack of compatible antibodies. However, we used a proximity ligation assay to demonstrate heterodimerization of cytoglobin with HMGB2. Immunofluorescence staining for heterodimers was evident in both the cytosol and nucleus of HEK293 cells expressing cytoglobin (**Fig. 6c**) and in sub-cultured human coronary smooth muscle cells (**Supplementary Fig. 7**).

**Figure 6.**
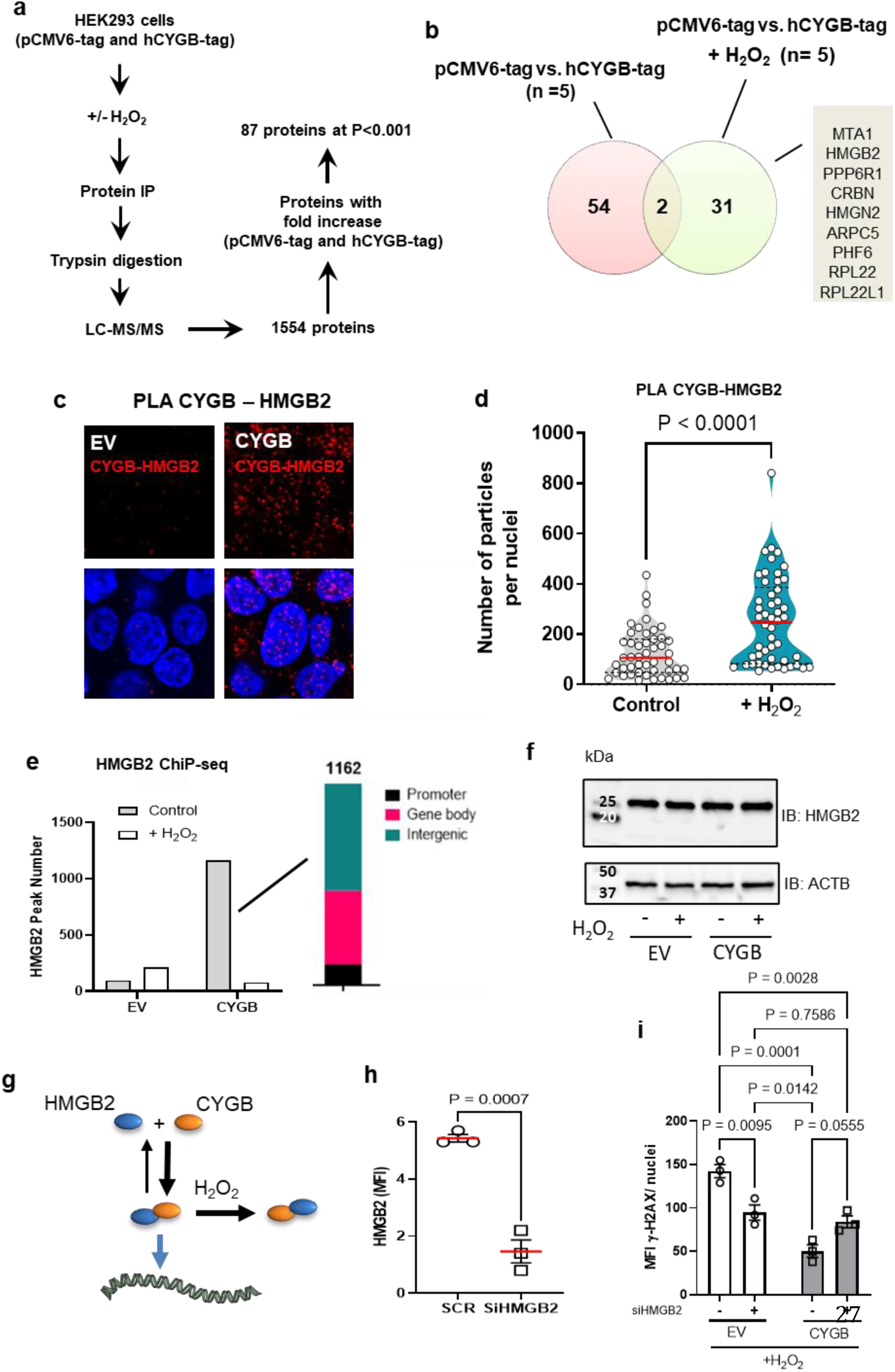
Cytoglobin regulates HMGB2 genome wide binding. (**a**) Schematic of the immunoprecipitation and mass spectrometry approach used to identify cytoglobin-interacting proteins in HEK293 cells stably expressing empty (pCMV6-tag) and hCYGB-tag vectors. **(b)** Venn diagram showing limited overlap of cytoglobin protein interactors between control and hydrogen peroxide treated samples. 54 proteins were interacting with CYGB before hydrogen peroxide addition and 31 proteins after hydrogen peroxide treatment. Subsequent analysis yielded several cytoglobin nuclear-associated proteins obtained from Co-IP MS/MS in hydrogen peroxide treated conditions. **(c)** Representative immunofluorescence images of proximity ligation assay for CYGB and HMGB2 in HEK293 cells with or without stable expression of cytoglobin (hCYGB); scale bar, 30 µm. **(d)** Quantitation of proximity ligation assay signal (number of nuclear particles per nuclei) of HEK293 cells stabling expressing empty vector (pCDNA) and human cytoglobin (hCYGB) following hydrogen peroxide treatment for 10 minutes. **(e)** HMGB2 ChiP-seq in pCDNA and hCYGB HEK 293 cells with or without treatment with hydrogen peroxide showing increase in HGMB2-genome wide binding in the presence of cytoglobin and loss of binding after treatment with hydrogen peroxide. Right panel: Peak position (in % peaks) of the ChiP analysis of HMGB2 **(f)** Western blot analysis of lysates from pCDNA (empty vector) and human cytoglobin (hCYGB) with and without hydrogen peroxide treatment probed for HMGB2 (24 kDa and β-Actin (ACTB, loading control, 40 kDa). **(g)** Cytoglobin heterodimerization is associated with an increase in HMGB2 genome-wide binding. Hydrogen peroxide (H2O2) increases cytoglobin-HMGB2 heterodimerization but inhibits HMGB2 genome-wide binding. **(h)** Quantitative analysis of HMGB2 mean fluorescence intensity (MFI) following HMGB2 silencing with siRNA targeting HMGB2 (SiHMGB2) or scrambled siRNAs (SCR); data are presented as +/-SEM; p values were determined by unpaired Student’s t-test. **(i)** HMGB2 was silenced in pCDNA (empty vector) and hCYGB (CYGB expressing) HEK cells, treated with 200 μM bolus hydrogen peroxide, and quantitated for %cells positive with γ-H2AX. Data are presented as +/-SEM; p values were determined by two-way ANOVA followed by Tukey’s post hoc test for multiple comparisons.

### Cytoglobin regulates HMGB2 genome wide binding and HMGB2-dependent activation of DNA repair pathways

As shown in **Fig. 6c**, we detected heterodimerization of cytoglobin with HMGB2 in HEK293 cells expressing cytoglobin, yet the protein-protein interaction analysis revealed heterodimerization only in the presence of hydrogen peroxide (**Fig. 6b and Supplementary Table 1 and 2**). Upon further analysis, we found an increase in cytoglobin and HMGB2 heterodimerization in the nucleus of HEK293 cytoglobin expressing cells after treatment with hydrogen peroxide (**Fig. 6d),** consistent with preferable pull-down following hydrogen peroxide treatment. To explore the functional significance of cytoglobin interaction with HMGB2, we performed chromatin immunoprecipitation of HMGB2 using a dual-crosslinking method followed by next-generation sequencing (ChIP-Seq)^28^. We identified only 93 HMGB2 peaks in HEK293 cells expressing the control empty vector (**Fig. 6e**). However, the expression of cytoglobin was associated with the detection of 1162 HMGB2 peaks, mostly bound to intergenic and gene body regions (**Fig. 6e**). We found that hydrogen peroxide treatment reversed this effect leading to peak numbers comparable to control (empty vector) conditions. Neither cytoglobin expression nor hydrogen peroxide treatment changed total HMGB2 protein content as determined by Western blot analysis (**Fig. 6f**). Overall, these results were consistent with a model where cytoglobin heterodimerization with HMGB2 increases genome wide binding of HMGB2 and hydrogen peroxide promotes the sequestration of cytoglobin –HMGB2 heterodimers away from HMGB2 DNA binding (**Fig. 6g**).

Past studies indicated that HMGB2 is necessary to initiate DNA repair, in part based on the observation that decrease HMGB2 expression followed by a genotoxic insult was associated with a decrease in γ-H2AX^29^. We first established that we could silence HMGB2 via siRNA in HEK293 cells that expressed the empty vector and hCYGB plasmid (**Fig. 6h**). We then confirmed that silencing of HMGB2 in the empty vector cells followed by hydrogen peroxide treatment decreased γH2AX when compared to cells transfected with scrambled control siRNA (**Fig. 6i**). Importantly, HMGB2 deletion was able to decrease γ-H2AX staining even in the presence of cytoglobin (**Fig. 6i**). These results suggest that cytoglobin interacts with HMGB2 to facilitate its removal from chromatin binding sites, which is required for functional DNA repair to inhibit hydrogen peroxide induced DNA damage.

### Cytoglobin heterodimerizes with HMGB2 in human intimal hyperplasia

To examine the possible significance of cytoglobin expression in human vessels, we immuno-stained for cytoglobin and HMGB2 in autopsy specimens of human temporal arteries. There was significant staining for both cytoglobin and HMGB2 in the medial and intimal regions of the vessels (**Fig. 7a**). We also performed a proximity ligation assay for cytoglobin and HMGB2 and established heterodimerizations of the two proteins within the nucleus of cells in the medial region of these vessels (**Fig. 7b**). These results are consistent with our studies performed *in vivo* in the mouse ligation model, and in vitro in sub-cultured human cells.

**Figure 7.**
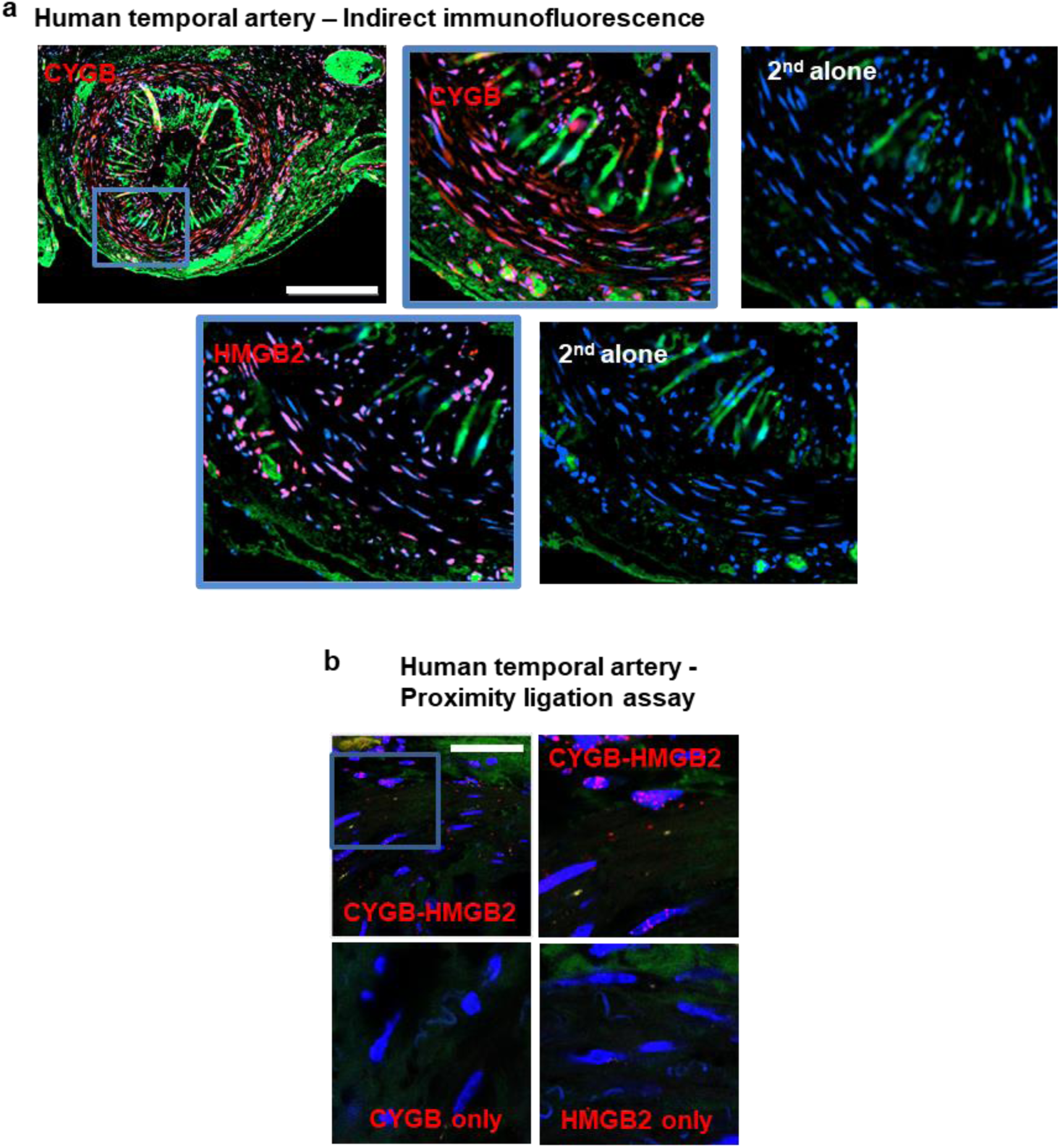
Cytoglobin and HMGB2 heterodimerizes in human vessels. (**a)** Human temporal artery samples were obtained, and indirect confocal immunofluorescence was performed for CYGB and HMGB2. Bar scale, 300 µm. **(b**) Proximity ligation assay for CYGB and HMGB2 from human temporal arteries; scale bar, 50 µm.

## Discussion

Despite extensive biochemical characterization over the past twenty years, the functional significance of cytoglobin is still unclear. In the present work, we reveal an unexpected mechanism by which cytoglobin regulates gene expression and DNA repair pathways in the vasculature. We combined transcriptional and protein profiling with functional validation *in vitro* and *in vivo* to establish a model where cytoglobin is trafficked into the nucleus, heterodimerizes with the non-histone DNA interacting protein HMGB2, and regulates HMGB2 genome-wide binding. This study extends the function of mammalian globins beyond oxygen transport and nitric oxide handling to direct regulation of gene expression and DNA repair pathways in the vasculature.

An important finding is that injury-induced suppression of SMC differentiation markers was accelerated in the carotid arteries of cytoglobin-deficient mice. The transcriptomic analysis performed in cultured human aortic vascular smooth muscle cells was also consistent with a role for cytoglobin in regulating smooth muscle gene programs. Contractile features of vascular smooth muscle cells are maintained by the concerted action of the transcription factor Serum Response Factor (SRF) and cofactors such as myocardin that restrain SRF activity to smooth muscle specific genes. Importantly, we show a decrease in myocardin mRNA transcripts, suggesting that cytoglobin regulates the expression of smooth muscle contractile genes through changes in myocardin levels. Previously, we showed that the loss of cytoglobin inhibits neointimal hyperplasia in the rat carotid angioplasty and mouse carotid ligation models^9^. While we did not look at mechanisms in the mouse model, we found that silencing of cytoglobin in the rat model was associated with increased apoptosis and cellular loss. Other studies have also indicated a role for cytoglobin in regulating cell fate through regulation of cell survival^9, 11, 12, 30, 31^. However, in the present work, we found no difference in medial and adventitial cell numbers between the wild-type and cytoglobin knockout mice, suggesting that increased cell death was not a primary contributor at least early in this injury model. Overall, our results indicate complex gene reprogramming. For example, we established that loss cytoglobin was also accompanied with activation of DNA repair pathways *in vitro* following oxidant treatment and *in vivo* following ligation. Thus, even if cytoglobin may accelerate smooth muscle cell de-differentiation, initiation of DNA damage might promote inhibition of clonal expansion of smooth muscle cells, leading to the absence of neointima in this model.

An open question when we began these studies was the molecular mechanisms by which cytoglobin may alter gene expression and DNA damage. To provide resolution, we first addressed the issue of cytoglobin subcellular localization. Due to its small size (∼21 kDa), it has been proposed that cytoglobin might diffuse passively between the cytosol and nucleus^20, 32^ while other studies suggested regulated accumulation through unknown processes^11^. In the present work, we show that the subcellular localization of cytoglobin in human aortic smooth muscle cells is regulated by growth factors as serum-starvation sequesters cytoglobin in the cytoplasm and serum-stimulation stimulates its nuclear accumulation. We also found that the nucleocytoplasmic trafficking of cytoglobin in response to serum stimulation is regulated by the karyopherins: importin α/β1 and exportin 1 (XPO1 or CRM1). Our results support a model where cytoglobin exclusion from the nucleus in quiescent smooth muscle cells is maintained through rapid CRM1-dependent nuclear export. Following growth-media stimulation, nuclear entry through importin-α/β dominates over nuclear export. While we did not screen for specific growth factors that might stimulate cytoglobin nuclear accumulation, we were intrigued with the possibility that intracellular redox signals might drive this process. We found that the nucleocytoplasmic shuttling of cytoglobin is an oxidant sensitive process. The antioxidant, N-acetyl cysteine inhibited the nuclear import, whereas the oxidant, hydrogen peroxide induced the nuclear import of cytoglobin. Significantly, we determined the expression of the hydrogen peroxide producing enzyme NOX4 was required for cytoglobin nuclear translocation. NOX4 has been shown to regulate many aspects of smooth muscle function including cell migration and differentiation. Thus, NOX4 regulates Serum Responsive Factor (SRF) protein levels and myocardin related transcription factor A (MRTFA) phosphorylation^33, 34^ – two important transcriptional regulators of smooth muscle differentiation^35, 36^. Interestingly, a recent study linked NOX4-derived hydrogen peroxide to endothelial cell differentiation through NOX4-dependent nuclear translocation of the histone demethylase Jumonji domain-containing protein-3 (JmjD3)^37^. In other instances, the molecular mechanism by which hydrogen peroxide regulates the nucleocytoplasmic shuttling of proteins has been ascribed to the redox regulated binding of the karyopherin CRM1 with its cargo protein^38^. Thus, considering the inhibition of cytoglobin nuclear accumulation following NOX4 silencing or pharmacological inhibition of CRM1, it is possible that a similar mechanism is at play. Cytoglobin possesses two surface cysteine residues that are oxidized by hydrogen peroxide to form a disulfide bridge and affects the binding properties and chemistry of cytoglobin’s heme center^39, 40^. Consequently, NOX4-dependent oxidation of these cysteine residues might be required to reconfigure cytoglobin structure and protein-protein interactions for effective nuclear import/export.

Although we obtained clear evidence of regulated nuclear accumulation of cytoglobin, this did not establish that nuclear cytoglobin was required to inhibit hydrogen peroxide induced DNA damage and regulate gene expression. This is not trivial since we recently found that cytoglobin scavenges intracellular hydrogen peroxide through redox cycling of its two cysteine residues^10^. Thus, in this case even cytosolic cytoglobin might be sufficient to limit the diffusion of hydrogen peroxide into the nucleus and inhibit the reaction of this oxidant with DNA. To answer this specific question, we leveraged our observation that the C-and N-terminal end truncated cytoglobin distinctly sequestered in the cytoplasm and nucleus, respectively. The C-truncated cytosolic cytoglobin was less effective at inhibiting DNA damage than the full length cytoglobin or the nuclear associated N-terminal truncated form. In fact, we showed that forced accumulation of the C-terminal truncated cytoglobin through fusion with a canonical nuclear localization sequence inhibited DNA damage. Surprisingly, we were unable to demonstrate that it could directly scavenge hydrogen peroxide in the nucleus based on changes in the hydrogen peroxide sensor Hyper-7 targeted to the nucleus. Moreover, the nuclear associated C-terminal truncated cytoglobin inhibited DNA damage induced by camptothecin, a topoisomerase I inhibitor that promotes DNA damage independent of hydrogen peroxide^25^. This led us to conclude that the inhibitory effect of cytoglobin on DNA damage cannot be explained solely based on its antioxidant activity.

We then sought to delineate a mechanism of cytoglobin function independent of its antioxidant capacity. Therefore, we identified a subset of nuclear protein interactors for cytoglobin that included high mobility group box 2 (HMGB2), metastasis-associated 1 (MTA1), serine/threonine-protein phosphatase 6 regulatory subunit 1 (PP6R1) and high mobility group nucleosomal binding domain 2 (HMGN2). Most significantly, all these interactors regulate chromatin remodeling and maintain genomic stability^26, 27, 41, 42^. Canonical High Mobility Group (HMG) proteins (including HMGB2 and HMGN2 identified in this study) bind DNA with limited specificity and exert twisting and bending structural constraints on DNA to promote or repress chromatin function^27^. Global reprogramming of chromatin accessibility by HMGB2 has been demonstrated^28, 43–46^ and chromatin compaction has also been shown to shield DNA strands from chemical damage^47^. We show that the inhibitory effect of cytoglobin on DNA damage depends on HMGB2. Most significantly, we were able to demonstrate that cytoglobin increased HMGB2 binding genome wide. This was lost following treatment of cells with hydrogen peroxide, although cytoglobin-HMGB2 heterodimerization was increased. These results are consistent with a model where cytoglobin stimulates HMGB2 binding to DNA cognate sequences to regulate chromatin accessibility and gene expression.

We demonstrated heterodimerization between cytoglobin and HMGB2, in smooth muscle cells *in vivo* and *in vitro*. In other settings, HMGB1 and 2 bind nuclear regulators of cell fate such as OCT4 or directly bind to specific promoter regions^46^. Studies by Lee and co-workers have shown that HMG-I, which belongs to the HMGA family, is expressed in smooth muscle cells^48^ and HMG-I/Y binds to and enhances serum responsive factor (SRF)-dependent activation of SM22α^49^. In contrast, Zhou and co-workers found that the protein HMG2-like1 (HMG2L1) binds to myocardin and inhibits myocardin-SRF complex binding to the promoters of smooth muscle specific genes^50^. Thus, we propose that the translocation of cytoglobin to the nucleus and the formation of the cytoglobin-HMGB2 heterodimer in vascular smooth muscle cells represent a coordinated response that is protective against DNA damage and regulates smooth muscle contractile gene expression. The literature outlined above would favor a model in which the cytoglobin-HMGB2 axis regulates gene transcription through specific interactions with transcriptional regulators such as myocardin. An alternative explanation would rely on a more promiscuous mechanism that controls higher order chromatin structure to coordinate large scale changes in transcriptional activity, like recent findings in cardiac remodeling^43, 51^ or vascular senescence^28^. While the transcriptional and epigenomic machinery that regulates smooth muscle gene programs has been relatively well characterized^36^, the impact of genome-wide changes in chromatin compaction and accessibility on specific gene programs associated with smooth muscle function is not understood.

In summary, our results challenge the current dogma related to mammalian globin functions simply as oxygen transport and nitric oxide handling systems. We establish a novel role for cytoglobin in regulating gene transcription and DNA repair pathways. Critical challenges for future studies will be to refine its mechanism of action related to vascular smooth muscle plasticity. Biochemical studies have detailed a unique redox chemistry at the heme center and through surface cysteine residues. It will be important to determine how this regulates cytoglobin nucleocytoplasmic shuttling and its interaction with DNA modifiers such as HMGB2. One exciting possibility is that the binding of small gaseous molecules such as molecular oxygen or nitric oxide to cytoglobin’s heme center might provide a means by which global changes in gene transcription required for smooth muscle phenotypic adaptation can be directly fine-tuned in response to environmental changes. Thus, it will be important to define the role of cytoglobin in smooth muscle cells in the context of different cardiovascular diseases such as atherosclerosis in which both environmental and genetic contributions are critical. We expect that these studies will provide new mechanistic insights that will contribute to the development of new therapeutics for cardiovascular diseases.

## Methods

### Supplies and reagents

All supplies and reagents are listed in Supplementary Table 1.

### Mice

All experiments involving mice were approved by the Institutional Animal Care and Use Committee at Albany Medical College. The cytoglobin “knock out first” allele was obtained from the University of Toronto, Canada, and global wild-type and cytoglobin knockout mice littermates (Cygb WT mice and Cygb KO) were generated from heterozygous breeding pairs. Both males and females were used and randomly assigned to experimental groups and all experiments were performed in 6-to 8-week-old littermates. For the carotid ligation studies, mice were anesthetized with inhaled isoflurane. After midline incision of the neck, complete ligation of the left common carotid artery was performed, just proximal to the carotid bifurcation. Carotids were harvested at the indicated time and processed for bulk transcriptomic analysis, Western blot, and immunofluorescence analysis.

### Transcriptomic analysis

Mouse carotid arteries from 3-day ligation experiments were homogenized in Trizol reagent using a mini bead mill homogenizer with a mixture of 1.4– and 2.8-mm Zirconium oxide beads (VWR). RNA was isolated according to the Tizol protocol and quantified on a Qubit Flex Fluorometer using the Qubit RNA HS assay kit. Library preparation was performed using an Ion Chef System followed by sequencing using an Ion GeneStudio S5 Plus System (All from Thermo Fisher Scientific, San Jose, CA). We used the Ion AmpliSeq Transcriptome Mouse Gene Expression Kit according to manufactures directions

### Generation of HEK 293 clones, cell culture, and treatments

Human cytoglobin was stably expressed in HEK293 cells as previously described using either pCDNA 3.1 plasmids or pCMV6 DYKDDDDK tagged plasmids^9, 10^. Briefly, stably expressing human cytoglobin cell lines were established through limiting dilution cloning. Cells were maintained under selection with Geneticin in DMEM supplemented with 10% fetal calf serum, 2 mM L-glutamine. For some experiments, HEK293 cells were treated with 50 µM camptothecin for 1 hour. For immunofluorescent staining clones were plated on poly-L-lysine treated 8 well glass Ibidi chamber slides with DMEM, 10% FBS, 2 mM L-glutamine. Cells were given 24 hours to attach, for experiments requiring serum starvation the media was removed, cells were rinsed with 37 °C phosphate buffered saline and incubated in DMEM with 0.2% FBS and 2 mM L-glutamine for 5 hrs.

Human Aortic and Coronary Smooth Muscle Cells were cultured as directed in VascuLife Smooth Muscle Cell Growth Media. (VascuLife Basal Media supplemented with VascuLife Smooth Muscle Cell Lifefactors kit (fetal bovine serum 5% v/v, basic human fibroblast growth factor (rhFGF) 5 ng/ml, human epidermal growth factor (rhEGF) 5 ng/ml, recombinant human insulin growth factor 5 µg/ml and Gentamicin (30µg/ml)/Amphotericin B (15 ng/ml) Cells were used between passages 4 and 10. Starvation media was prepared with VascuLife Basal media and 0.2% FBS.

For all experiments, human vascular smooth muscle cells were plated in growth media for 24 hours. For Ivermectin B treatment, the cells were placed in starvation media for 5 hrs then 25 ng/mL Ivermectin in growth media for 12 hrs. The Leptomycin B treated cells were incubated in starvation media containing 20 ng/ml Leptomycin B for 12 hours. The hydrogen peroxide treated cells were serum starved for 5 hrs, then treated with 10-200 µM Hydrogen Peroxide in starvation media for varying times. The N-acetyl Cysteine or DPI treated cells were serum starved for 5 hours then incubated with either 1 mM N-acetyl cysteine or 10 µM diphenyleneiodonium (DPI) prepared in growth media overnight. For immunofluorescence studies, human smooth muscle cells were plated in 6 well 1.5μ Ibidi slides. For experiments including serum starvation, the growth media was replaced with 0.2% starvation media for 5 hours.

### Electroporation and Transfection

The silencing of Cytoglobin, NOX4 and NOX5 in smooth muscle cells was achieved by nucleofection using program U025 on the Amaxa Nucleofector II. Each reaction introduced 100 nM siRNA into 1^6 cells following manufacture instructions for the Amaxa Human Aortic Smooth muscle cells Nucleofector kit. Cells were silenced for 72 hrs prior to experimentation. Transfection of siHMGB2 into the HEK293 clones was done according to the manufacturer’s instructions for Dharmafect 1. 50 nM of siRNA was introduced into cells pre-plated in DMEM, 10% FBS and 2 mM L-glutamine, and allowed to incubate for 72 hrs.

### Microarray Analysis

We profiled the expression of transcripts in human aortic vascular smooth muscle cells using the Clariom D human DNA microarray (Thermo Fisher Scientific Inc.). This platform includes information from probe sets representing known and predicted exons and introns on both strands of the genome that have been mapped to more than genes. The experimental design consisted of three groups of human aortic vascular smooth muscle cells: control, and electroporated with either scrambled siRNA or human cytoglobin siRNA, each in triplicate. Intensity data was log-2 transformed and normalized using the Transcriptomics Analysis Console from Affymetrix. We used a fold change ≤ –1.5 or ≥ 1.5 and a P value of ≤ 0.05 to filter transcripts that were differentially expressed in the siCygb group compared to untreated. From this list, transcripts that were found to be statistically different between the untreated and the scrambled siRNA group were removed. Post-analysis of the differentially expressed gene set was performed using Ingenuity Pathway Analysis (QIAGEN Inc.) ^52^.

### Chromatin Immunoprecipitation (ChIP)

HEK Cells expressing either EV or Human Cytoglobin were incubated with and without 150 µM hydrogen peroxide for 10 minutes. We utilized the SimpleChIP Enzymatic Chromatin IP kit from Cell Signaling following the manufactures protocol with a few exceptions. Cells were crosslinked 1% PFA for 1 hour at 4°C. For each 10 µg of digested Chromatin, we used 7.5 µg of the immunoprecipitating HMGB2 antibody. The purified DNA was sent for next generation sequencing (ChIPSeq).

### Real-time PCR

RNA was extracted from cells by direct lysis in Trizol Reagent. Total RNA was isolated according to manufactures directions. cDNA was synthesized using Qiagen’s QuantiTect Reverse Transcription Kits. qPCR analysis was conducted using gene-specific primers, and SsoAdvanced Universal SYBR Green super mix on an BioRad CFX Connect Realtime System equipped with CFX Maestro software. Oligonucleotide primers were designed using PrimerBLAST (NCBI) and purchased from IDT (Coralville, IA).

### Protein extraction and Western blotting

Western blotting was performed as previously described^9^. Briefly protein lysates were prepared from cultured cells scraped directly into RIPA buffer with HALT Protease and Phosphatase Inhibitor Cocktail. Protein concentrations were determined using a BCA Protein assay. Lysates were then heat denatured with either 2 or 4X Laemmli sample buffer, equal protein loads were added to corresponding wells on 4-20% gradient gels. After separation the proteins were transferred onto 0.2 µM PVDF membranes and blocked in Tris-buffered saline containing 0.1% Tween-20 and 5% nonfat milk (TBST) for 1 hr at RT. The membranes were then probed with primary antibodies for 1 hr at RT or overnight at 4°C, washed 3X in TBST then incubated with the corresponding HRP conjugated secondary antibodies. Signals were detected with Clarity Western ECL substrate on a BioRad ChemiDoc MP Imaging System equipped with Image lab software. The primary and secondary antibodies are listed in Supplementary Table 1.

### Immunoprecipitation

Cells lysates from both untreated HEK Cells stably transfected with empty pCMV6 or human cytoglobin pCMV6 DYKDDDDK tagged vectors or those treated with 150 uM hydrogen peroxide for 10– minutes were collected in cold IgG lysis buffer with 1:50 HALT protease and phosphatase inhibitor. The lysates were sheard through progressively smaller gauge needles, cleared by centrifugation at 12 000 × g for 20 minutes at 4°C. The IP was performed with Pierce Anti-DYKDDDDK Magnetic Agarose beads according to manufactures protocol. Briefly, 800 ug of protein lysate, diluted to a final volume of 300 µl with lysis buffer, was added to 100 µl of magnetic agarose slurry, tubes were rotated at room temperature for 20 minutes, beads were washed 3 times with PBS and once with ultrapure water before being eluted in 100 µl of 8 M urea, 100 mM Tris (pH-7.5) and 100 mM NaCl

### Mass spectrometry

The 100 µl Immunoprecipitation eluates were spiked with ∼10 µl of a 10x stock solution of TCEP and chloroacetamide in 50 mM Tris (pH∼8) to achieve a final concentration of 10 and 40 mM, respectively. The mixtures were vortexed for 10 min, and 1.5 µl of MS grade trypsin was added to each sample. The samples were digested overnight at ambient temperature on a rocker, an additional 1 µl of trypsin was added to each sample in the morning. The samples were then incubated for 2 additional hours, acidified by addition of 10% TFA to the final concentration of 1%, and desalted using StrataX SPE cartridges (Phenomenex).

For data acquisition, each sample was resuspended in 15 µl 0.2% formic acid. 2.2 µl of each sample were injected on a 75 µm ID × 360 µm OD column packed in-house^53^ with 1.7 um BEH C18 particles (Waters) to the final length of 35 cm. The column was kept at 50°C, and the peptides were separated over a 60 min gradient using Dionex nanoUltiMate 3000 chromatographic system at flow rate of 315 nl/min. The peptides were ionized at 2.2 kV and introduced into Orbitrap Eclipse (Thermo Scientific) for analysis. MS1 scans were obtained in the Orbitrap at resolution of 60,000, AGC target of 200%, a maximum injection time (max IT) of 50 ms, and over the range of 300-1350 m/z. The precursor ions with charge states of 2-5 were fragmented at normalized collision energy of 25%, and subsequent MS2 analysis was performed in the Orbitrap at resolution of 7500 with max IT of 25 ms with “Define 1^st^ mass” set to 150. Dynamic exclusion was set to 15 s and the quadrupole isolation width was set to 0.7 Th.

For data processing, RAW files were searched in MaxQuant (version 1.6.10.43)^54^, against a database of canonical proteins and isoforms (*Homo sapiens*, downloaded from Uniprot 20190619). All parameters were set to defaults, except for “digestion mode” was set to Trypsin (no proline), LFQ quantification was enabled with the count of 2, and match-between-runs was enabled. The final protein group results were filtered for reverse identifications, contaminants, and “identified by site only.” Proteins groups containing missing values in more than 50% of samples were removed. The remaining missing values were input using the web-based tool Argonaut^55^.

Raw data have been deposited to the MassIVE database under the accession number MSV000091289. The files can be accessed using the log-in credentials: Username: MSV000091194_reviewer; Password: Cytoglobin.

### Immunofluorescence

Mouse carotid tissues, 10 µm sections were cut from prepared OCT blocks using a Leica CM1850 cryostat and transferred to charged microscope slides, allowed to dry at room temperature and stored at –80°C until use. Slides were removed from the freezer, air dried, fixed with ice cold acetone for 10 minutes, air dried again, then outlined with a hydrophobic barrier pen. Tissues were briefly rehydrated with PBS, blocked with RTU animal free blocker and diluent for at least 1 hour. Primary antibodies or corresponding isotype controls, diluted in blocking/diluent buffer, were applied to the samples for one hour at room temperature or overnight at 4°C. Slides were washed 3X in PBS/0.1% Triton X prior to a one-hour incubation with the secondary antibodies. Slides were washed once with PBS/0.1% Triton X and once with a 50:50 PBS/water mix; incubated with 1 µM DAPI for 15 minutes at room temperature then rinsed again in the 50:50 PBS/water mixture; cover slipped with VectaShield Antifade Mounting Medium; sealed with clear nail polish and stored protected from light at 4°C.

HEK and Human Aortic Smooth Muscle Cells; After removal of growth media, cells were immediately fixed in room temperature 4% paraformaldehyde for 15 minutes. Slides were rinsed with PBS and cells were permeabilized for 5 minutes in PBS/0.2% Triton-X100, blocked with 5% sera representing the secondary antibody species for at least 1 hour. Cells were incubated with primary antibodies or matched isotype controls diluted in blocking buffer for 1 hour at RT, washed 3 × 5 minutes in PBS/0.1% Triton X 100, incubated with corresponding secondary antibody for 45 min at RT, washed once in PBS/0.1% Triton X 100, once in 50:50 PBS/water mix then stained with 1 µM DAPI for 15 minutes at room temperature. Slides were rinsed in the 50:50 PBS/water mixture; covered slipped with VectaShield Antifade Mounting Medium (8 well removable chamber) or sealed with Ibidi mounting media (1.5μ Ibidi 6 chamber slides).

The mean fluorescence intensity (MFI) ratio of nuclear *vs.* cytoplasmic per cell was determined and quantified using the Intensity Ratio Nuclei Cytoplasm Tool from FIJI ImageJ software and plotted as MFI (nucleus/cytoplasm). Fiji (ImageJ) software (National Institutes of Health, Bethesda, MD, USA) was used for analysis and quantification of images.

### Proximity Ligation Assay

All steps performed in a 37°C humidity chamber. After fixation or fixation and permeabilization as described above, proximity ligation assays were performed according to Sigma’s Duolink PLA Fluorescence protocol. Briefly, after blocking the samples were incubated with primary antibodies. After gentle washing in wash buffer A the plus (rabbit) and minus (mouse) PLA probes were added to samples and incubated for 1 hr, slides were washed 2 × 5 minutes with wash buffer A then incubated with ligation solution for 30 minutes. Slides were washed again in Buffer A 2 × 2 minutes. Samples were incubated with amplification solution for 100 minutes. Samples were washed in Wash buffer B 2 × 10 minutes, incubated with 1 µM DAPI for 15 minutes and rinsed once more in 0.1% Buffer B prior to being cover slipped with VectaShield Antifade Mounting Medium.

### Real-time imaging

For imaging experiments, HEK 293 cells with a stable transfection of human cytoglobin or empty vector were seeded onto Poly-L-lysine coated 35 mm glass bottom dishes with DMEM, 10% FCS, 2 mM L-Glutamine, no antibiotic. After 24 hours, cells were transfected with the Hyper 7 plasmids using Dharmafect kb transfection reagent according to the manufacturer’s recommendations. Twenty-four hours after transfection, culture medium was replaced with 1.2 mL of HBSS supplemented with 20 mM HEPES. Cell imaging was performed using a Leica DMI 8 Thunder microscope, equipped with a HC PL APO 63x 1.4NA oil objective at 37 °C. Samples were excited sequentially via 440/15 and 510/15 band-pass excitation filters. Emission was collected every 20 seconds using a 519/25 bandpass emission filter. After 5–10 images were acquired, a small volume of hydrogen peroxide was carefully added to obtain a final concentration of 200 µM. The time series were analyzed via Fiji freeware (https://fiji.sc). The background was subtracted from 440 and 510 nm stacks. Every 510 nm stack was divided by the corresponding 440 nm stack frame-by-frame. The resulting stack was depicted in pseudo-colors using a “16– colors” lookup table. The plasmid pCS2+Hyper7-NLS was a gift from Dr. Belousov (Addgene plasmid # 136468).

### COMET Assay

The comet assay was performed with the R&D Comet assay kit according to the manufacturer’s instructions. Human smooth muscle and HEK293 cells were mixed into LMAgarose (at 37 °C) in a 1:10 vol ratio for 20 minutes and immobilized onto a comet assay glass slide (50 μL each well). The glass slides were placed in the dark at 4°C for 30 min. Following this step, slides were immersed in lysis solution at 4 °C overnight. The excess buffer was drained, and the slides were immersed in alkaline unwinding solution for 1 hr in the dark at 4 °C. Later, the slides were placed in a gel box with 850 mL alkaline electrophoresis solution at 4 °C then electrophoresed at 25 V for 30 min in the dark at 4°C. Excess electrophoresis solution was removed, and the slides were immersed twice in distilled H_2_O for 5 minutes each and then in 70% ethanol for 5 minutes. Slides were then dried at 37°C for 15 minutes. The nuclei were stained with 100 μL of DAPI at room temperature in the dark for 30 min. The fluorescent images were observed on Biotek Cytation5 imager. DNA damage and migration were assessed (100 cells in each group) using the GEN 5 software package.

### Human sample harvesting, tissue processing and immunofluorescence staining

Deidentified archival temporal artery specimens were obtained from the Department of Pathology & Laboratory (Institutional Review Board protocol IRB#). For immunostaining and proximity ligation assay, slides with FFPE sections were pretreated with Trilogy to deparaffinize, rehydrate and unmask immunogens according to manufactures instructions. The tissues were then ready for either antibody staining or proximity ligation assay according to methods listed above.

### Statistical analysis

Statistical analyses were performed with GraphPad Prism 9.0. The statistical test used to analyze each data set is specified in individual figure legends and p-values are shown in figures. A p-value less than 0.05 was considered statistically significant. Results are expressed as mean ± SEM, and statistical analysis using unpaired t-tests, one-or two-way ANOVA for two or two or more groups comparison followed by Tukey’s multiple comparison test were used. The number of independent replicates, and statistical tests used are indicated in the figure legends, when applicable.

## Acknowledgments

This work was supported by NIH grants RO1 HL142807 (to D.J.), P41 GM108538 (to J.J.C.), R01 CA233188 (to M.B.), KO1 HL130704 (to A.J.), and RO1 HL160661 (to A.J.).

## Author Contributions

Conceived and designed the analysis: C.M., F.J., R.I.L.S., and D.J. Collected data: C.M., F.J., L.G.C.P., K.G., D.H., J.B., S.V.C., J.B.C., E.S. Contributed data or analysis tools: D.J., A.J., S.V.C., M.R., R.P.W., I.A., S.J.S., M.B., J.J.C.; Performed the analysis; C.M., F.J., L.G.C.P., S.V.C., E.S. Wrote the paper: C.M., F.J., E.S., and D.J.

## Competing Interests

JJC is a consultant for Thermo Fisher Scientific, 908 Devices, and Seer.

## Literature Cited

1 Schechter, A. N. Hemoglobin research and the origins of molecular medicine. Blood 112, 3927–3938, doi:10.1182/blood-2008-04-078188 (2008).

2 Qiu, Y., Sutton, L. & Riggs, A. F. Identification of myoglobin in human smooth muscle. J Biol Chem 273, 23426–23432 (1998).

3 Straub, A. C., et al. Endothelial cell expression of haemoglobin alpha regulates nitric oxide signalling. Nature 491, 473–477, doi:10.1038/nature11626 (2012).

4 Halligan, K. E., Jourd’heuil, F. L. & Jourd’heuil, D. Cytoglobin is expressed in the vasculature and regulates cell respiration and proliferation via nitric oxide dioxygenation. J Biol Chem 284, 8539–8547, doi:10.1074/jbc.M808231200 (2009).

5 Smagghe, B. J., Trent, J. T., 3rd & Hargrove, M. S. NO dioxygenase activity in hemoglobins is ubiquitous in vitro, but limited by reduction in vivo. PloS one 3, e2039, doi:10.1371/journal.pone.0002039 (2008).

6 Amdahl, M. B., Sparacino-Watkins, C. E., Corti, P., Gladwin, M. T. & Tejero, J. Efficient Reduction of Vertebrate Cytoglobins by the Cytochrome b5/Cytochrome b5 Reductase/NADH System. Biochemistry 56, 3993–4004, doi:10.1021/acs.biochem.7b00224 (2017).

7 Zweier, J. & Ilangovan, G. Regulation of Nitric Oxide Metabolism and Vascular Tone by Cytoglobin. Antioxid Redox Signal, doi:10.1089/ars.2019.7881 (2019).

8 Liu, X., et al. Cytoglobin regulates blood pressure and vascular tone through nitric oxide metabolism in the vascular wall. Nat Commun 8, 14807, doi:10.1038/ncomms14807 (2017).

9 Jourd’heuil, F. L., et al. The Hemoglobin Homolog Cytoglobin in Smooth Muscle Inhibits Apoptosis and Regulates Vascular Remodeling. Arterioscler Thromb Vasc Biol, doi:10.1161/ATVBAHA.117.309410 (2017).

10. Mathai, C. et al. A role for cytoglobin in regulating intracellular hydrogen peroxide and redox signals in the vasculature. bioRxiv, 2023.2003.2031.535146, doi:10.1101/2023.03.31.535146 (2023).

11 Singh, S., et al. Cytoglobin modulates myogenic progenitor cell viability and muscle regeneration. Proc Natl Acad Sci U S A 111, E129–138, doi:10.1073/pnas.1314962111 (2014).

12 Zhang, S., et al. Cytoglobin Promotes Cardiac Progenitor Cell Survival against Oxidative Stress via the Upregulation of the NFkappaB/iNOS Signal Pathway and Nitric Oxide Production. Sci Rep 7, 10754, doi:10.1038/s41598-017-11342-6 (2017).

13 Thuy le, T. T., et al. Promotion of liver and lung tumorigenesis in DEN-treated cytoglobin-deficient mice. The American journal of pathology 179, 1050–1060, doi:10.1016/j.ajpath.2011.05.006 (2011).

14 Thuy le, T. T., et al. Absence of cytoglobin promotes multiple organ abnormalities in aged mice. Sci Rep 6, 24990, doi:10.1038/srep24990 (2016).

15 Hodges, N. J., Innocent, N., Dhanda, S. & Graham, M. Cellular protection from oxidative DNA damage by over-expression of the novel globin cytoglobin in vitro. Mutagenesis 23, 293–298, doi:10.1093/mutage/gen013 (2008).

16 Bennett, M. R., Sinha, S. & Owens, G. K. Vascular Smooth Muscle Cells in Atherosclerosis. Circ Res 118, 692–702, doi:10.1161/CIRCRESAHA.115.306361 (2016).

17 Herring, B. P., Hoggatt, A. M., Griffith, S. L., McClintick, J. N. & Gallagher, P. J. Inflammation and vascular smooth muscle cell dedifferentiation following carotid artery ligation. Physiol Genomics 49, 115–126, doi:10.1152/physiolgenomics.00095.2016 (2017).

18 Olivieri, M., et al. A Genetic Map of the Response to DNA Damage in Human Cells. Cell 182, 481–496 e421, doi:10.1016/j.cell.2020.05.040 (2020).

19 Ivashkevich, A., Redon, C. E., Nakamura, A. J., Martin, R. F. & Martin, O. A. Use of the gamma-H2AX assay to monitor DNA damage and repair in translational cancer research. Cancer Lett 327, 123–133, doi:10.1016/j.canlet.2011.12.025 (2012).

20. Geuens, E. et al. A globin in the nucleus! J Biol Chem 278, 30417-30420, doi:10.1074/jbc.C300203200 (2003).

21 Wagstaff, K. M., Sivakumaran, H., Heaton, S. M., Harrich, D. & Jans, D. A. Ivermectin is a specific inhibitor of importin alpha/beta-mediated nuclear import able to inhibit replication of HIV-1 and dengue virus. The Biochemical journal 443, 851–856, doi:10.1042/BJ20120150 (2012).

22 Nishi, K., et al. Leptomycin B targets a regulatory cascade of crm1, a fission yeast nuclear protein, involved in control of higher order chromosome structure and gene expression. J Biol Chem 269, 6320–6324 (1994).

23 Pak, V. V., et al. Ultrasensitive Genetically Encoded Indicator for Hydrogen Peroxide Identifies Roles for the Oxidant in Cell Migration and Mitochondrial Function. Cell Metab 31, 642–653 e646, doi:10.1016/j.cmet.2020.02.003 (2020).

24 de Cubas, L., Pak, V. V., Belousov, V. V., Ayte, J. & Hidalgo, E. The Mitochondria-to-Cytosol H(2)O(2) Gradient Is Caused by Peroxiredoxin-Dependent Cytosolic Scavenging. Antioxidants (Basel) 10, doi:10.3390/antiox10050731 (2021).

25 Hertzberg, R. P., Caranfa, M. J. & Hecht, S. M. On the mechanism of topoisomerase I inhibition by camptothecin: evidence for binding to an enzyme-DNA complex. Biochemistry 28, 4629–4638, doi:10.1021/bi00437a018 (1989).

26 Xue, Y., et al. NURD, a novel complex with both ATP-dependent chromatin-remodeling and histone deacetylase activities. Molecular cell 2, 851–861, doi:10.1016/s1097-2765(00)80299-3 (1998).

27 Stros, M. HMGB proteins: interactions with DNA and chromatin. Biochim Biophys Acta 1799, 101–113, doi:10.1016/j.bbagrm.2009.09.008 (2010).

28 Zirkel, A., et al. HMGB2 Loss upon Senescence Entry Disrupts Genomic Organization and Induces CTCF Clustering across Cell Types. Molecular cell 70, 730–744 e736, doi:10.1016/j.molcel.2018.03.030 (2018).

29 Krynetskaia, N. F., Phadke, M. S., Jadhav, S. H. & Krynetskiy, E. Y. Chromatin-associated proteins HMGB1/2 and PDIA3 trigger cellular response to chemotherapy-induced DNA damage. Mol Cancer Ther 8, 864–872, doi:10.1158/1535-7163.MCT-08-0695 (2009).

30 Latina, A., et al. DeltaNp63 targets cytoglobin to inhibit oxidative stress-induced apoptosis in keratinocytes and lung cancer. Oncogene 35, 1493–1503, doi:10.1038/onc.2015.222 (2016).

31 Tian, S. F., et al. Mechanisms of neuroprotection from hypoxia-ischemia (HI) brain injury by up-regulation of cytoglobin (CYGB) in a neonatal rat model. J Biol Chem 288, 15988–16003, doi:10.1074/jbc.M112.428789 (2013).

32 Schmidt, M., et al. Cytoglobin is a respiratory protein in connective tissue and neurons, which is up-regulated by hypoxia. J Biol Chem 279, 8063–8069, doi:10.1074/jbc.M310540200 (2004).

33 Clempus, R. E., et al. Nox4 is required for maintenance of the differentiated vascular smooth muscle cell phenotype. Arterioscler Thromb Vasc Biol 27, 42–48, doi:10.1161/01.ATV.0000251500.94478.18 (2007).

34 Lee, M., San Martin, A., Valdivia, A., Martin-Garrido, A. & Griendling, K. K. Redox-Sensitive Regulation of Myocardin-Related Transcription Factor (MRTF-A) Phosphorylation via Palladin in Vascular Smooth Muscle Cell Differentiation Marker Gene Expression. PloS one 11, e0153199, doi:10.1371/journal.pone.0153199 (2016).

35 Owens, G. K., Kumar, M. S. & Wamhoff, B. R. Molecular regulation of vascular smooth muscle cell differentiation in development and disease. Physiol Rev 84, 767–801, doi:10.1152/physrev.00041.2003 (2004).

36 Gomez, D., Swiatlowska, P. & Owens, G. K. Epigenetic Control of Smooth Muscle Cell Identity and Lineage Memory. Arterioscler Thromb Vasc Biol 35, 2508–2516, doi:10.1161/ATVBAHA.115.305044 (2015).

37 Hahner, F., et al. Nox4 promotes endothelial differentiation through chromatin remodeling. Redox Biol 55, 102381, doi:10.1016/j.redox.2022.102381 (2022).

38 Kwak, M. S., et al. Peroxiredoxin-mediated disulfide bond formation is required for nucleocytoplasmic translocation and secretion of HMGB1 in response to inflammatory stimuli. Redox Biol 24, 101203, doi:10.1016/j.redox.2019.101203 (2019).

39 Hamdane, D., et al. The redox state of the cell regulates the ligand binding affinity of human neuroglobin and cytoglobin. J Biol Chem 278, 51713–51721, doi:10.1074/jbc.M309396200 (2003).

40 Mathai, C., Jourd’heuil, F. L., Lopez-Soler, R. I. & Jourd’heuil, D. Emerging perspectives on cytoglobin, beyond NO dioxygenase and peroxidase. Redox Biol 32, 101468, doi:10.1016/j.redox.2020.101468 (2020).

41 Nanduri, R., et al. Epigenetic regulation of white adipose tissue plasticity and energy metabolism by nucleosome binding HMGN proteins. Nat Commun 13, 7303, doi:10.1038/s41467-022-34964-5 (2022).

42 Douglas, P., et al. Protein phosphatase 6 interacts with the DNA-dependent protein kinase catalytic subunit and dephosphorylates gamma-H2AX. Mol Cell Biol 30, 1368–1381, doi:10.1128/MCB.00741-09 (2010).

43 Monte, E., et al. Reciprocal Regulation of the Cardiac Epigenome by Chromatin Structural Proteins Hmgb and Ctcf: IMPLICATIONS FOR TRANSCRIPTIONAL REGULATION. J Biol Chem 291, 15428–15446, doi:10.1074/jbc.M116.719633 (2016).

44 Celona, B., et al. Substantial histone reduction modulates genomewide nucleosomal occupancy and global transcriptional output. PLoS Biol 9, e1001086, doi:10.1371/journal.pbio.1001086 (2011).

45 Aird, K. M., et al. HMGB2 orchestrates the chromatin landscape of senescence-associated secretory phenotype gene loci. J Cell Biol 215, 325–334, doi:10.1083/jcb.201608026 (2016).

46 Campbell, P. A. & Rudnicki, M. A. Oct4 interaction with Hmgb2 regulates Akt signaling and pluripotency. Stem Cells 31, 1107–1120, doi:10.1002/stem.1365 (2013).

47 Brambilla, F., et al. Nucleosomes effectively shield DNA from radiation damage in living cells. Nucleic Acids Res 48, 8993–9006, doi:10.1093/nar/gkaa613 (2020).

48 Pellacani, A., et al. Induction of high mobility group-I(Y) protein by endotoxin and interleukin-1beta in vascular smooth muscle cells. Role in activation of inducible nitric oxide synthase. J Biol Chem 274, 1525–1532, doi:10.1074/jbc.274.3.1525 (1999).

49 Chin, M. T., et al. Enhancement of serum-response factor-dependent transcription and DNA binding by the architectural transcription factor HMG-I(Y). J Biol Chem 273, 9755–9760, doi:10.1074/jbc.273.16.9755 (1998).

50 Zhou, J., Hu, G. & Wang, X. Repression of smooth muscle differentiation by a novel high mobility group box-containing protein, HMG2L1. J Biol Chem 285, 23177-23185, doi:10.1074/jbc.M110.109868 (2010).

51 Rosa-Garrido, M., et al. High-Resolution Mapping of Chromatin Conformation in Cardiac Myocytes Reveals Structural Remodeling of the Epigenome in Heart Failure. Circulation 136, 1613–1625, doi:10.1161/CIRCULATIONAHA.117.029430 (2017).

52 Kramer, A., Green, J., Pollard, J., Jr. & Tugendreich, S. Causal analysis approaches in Ingenuity Pathway Analysis. Bioinformatics 30, 523–530, doi:10.1093/bioinformatics/btt703 (2014).

53 Shishkova, E., Hebert, A. S., Westphall, M. S. & Coon, J. J. Ultra-High Pressure (>30,000 psi) Packing of Capillary Columns Enhancing Depth of Shotgun Proteomic Analyses. Anal Chem 90, 11503–11508, doi:10.1021/acs.analchem.8b02766 (2018).

54 Cox, J. & Mann, M. MaxQuant enables high peptide identification rates, individualized p.p.b.-range mass accuracies and proteome-wide protein quantification. Nat Biotechnol 26, 1367–1372, doi:10.1038/nbt.1511 (2008).

55 Brademan, D. R., et al. Argonaut: A Web Platform for Collaborative Multi-omic Data Visualization and Exploration. Patterns (N Y) 1, doi:10.1016/j.patter.2020.100122 (2020).

