## Supplementary Information for "Nuclear cytoglobin associates with HMGB2 and regulates DNA damage and genome-wide transcriptional output in the vasculature"

**SUPPLEMENTARY TABLE 1**

**SUPPLEMENTARY TABLE 2**

**SUPPLEMENTARY TABLE 3**

**SUPPLEMENTARY FIGURES**

### SUPPLEMENTARY TABLE 1

#### Supplies and Reagents

| Cells/siRNA/Plasmids | Supplier | Cat# |
|---|---|---|
| Cells HEK 293 | ATCC | CRL-1573 |
| Human Aortic Smooth muscle cells | Lifeline Cell Technologies | FC-0015 |
| Human Coronary Smooth muscle cells | Lifeline Cell Technologies | FC-0031 |
| ON-TARGETplus Human NOX4 siRNA | Horizon Discovery | L-0140194-00-0005 |
| ON-TARGETplus Human NOX5 siRNA | Horizon Discovery | L-010195-00-0005 |
| ON-TARGETplus Human CYGB siRNA | Horizon Discovery | L-016960-01-0010 |
| ON-TARGETplus Human HMGB2 siRNA | Horizon Discovery | L-011689-00-0005 |
| ON-Targetplus non targeting control pool (Scramble) | Horizon Discovery | D-001810-05 |
| DharmaFect 1 (transfection reagent) | Horizon Discovery | T-2001-03 |
| DharmaFect kb (transfection reagent) | Horizon Discovery | T-2006-01 |
| Amaxa Human Aortic Smooth muscle cells Nucleofector kit | Lonza | VPC-1001 |
| pcDNA 3.1 (empty vector) | Genscript | on request |
| pcDNA 3.1-hCYGB | Genscript | OHu14114C |
| pcMV6 Tag (empty vector) | Origene | PS100001 |
| pcMV6 hCYGB Tag | Origene | RC206642 |
| pCS2+HyPer7 | Addgene | 136466 |
| pCS2+HyPer7-NES | Addgene | 136467 |
| pCS2+HyPer7-NLS | Addgene | 136468 |

| Reagents | | |
|---|---|---|
| 37% formaldehyde solution | Sigma | 252549-100 |
| Acetone | Sigma | 179124-1L |
| Anti-DKYDDDDK Magnetic Agarose | Thermo Scientific | A36797 |
| BCA Protein Assay kit (Pierce) | Thermo Scientific | 23225 |
| βME (2-Mercaptoethanol) | Simga | 63689 |
| Camptothecin | Enzo | ALX-350-015-MO50 |
| Clairty Western ECL Substrate | BioRad | 1705061 |
| Diamidino 2 phenyl indole (DAPI) | Sigma | D9542-5mg |
| Diphenyleneiodonium chloride (DPI) | Sigma | D2926-10MG |
| DMEM (high Glucose) | Corning | 10-031-CV |
| DPBS | Corning | 21-031-CV |
| FBS | Cytiva | SH30396-03 |
| Geneticin (G418 sulfate) | Gibco | 10131-035 |
| HALT protease/phosphatase inhibitor | Thermo Scientific | 1861284 |

| | | |
|---|---|---|
| HBSS | Corning | 21-031-CV |
| HBSS with calcium and magnesium | Corning | 21-023-CV |
| HEPES | Sigma | H4034-100 |
| Hydrogen Peroxide | Sigma | 216763 |
| IP Lysis buffer - Pierce | Thermo Scientific | 87788 |
| Ivermection | Sigma | I8898 |
| Laemmli Sample buffer 2X | BioRad | 161-0737 |
| Laemmli Sample buffer 4X | BioRad | 161-0704 |
| Leptomycin B | Sigma | L2913 |
| L-Glutamine 200 mM | Sigma | G7513 |
| Molecular weigh markers (Percision Plus Western C) | BioRad | 161-0385 |
| N-acetyl cysteine | Sigma | A9165 |
| NaCl | Sigma | S7653 |
| Non-Fat Milk powder | Fisher | 50-751-7665 |
| OCT compound | Tissue-Tek | 4583 |
| Optimem | Gibco | 31985-062 |
| Poly-L-lysine solution | Sigma | P4832 |
| RIPA buffer | Sigma | R0278 |
| SsoAdvanced Universal Sybr Green Supermix | BioRad | 1725271 |
| Trilogy | Cell Marque | 920P-06 |
| Triton X-100 | Sigma | T9284 |
| Trizol Reagent | Life Technologies | 15596018 |
| Trypsin 0.25%, 2.21mM EDTA 1X | Corning | 25-053-CI |
| Trypsin-EDTA 0.5% no phenol red | Life technologies | 15400054 |
| Tween 20 | Sigma | P2287 |
| Urea | Sigma | U5378 |
| VascuLife Smooth Muscle Cell Basal Media | Lifeline Cell Technologies | LM-0002 |
| VascuLife Smooth Muscle Cell Lifefactors Kit | Lifeline Cell Technologies | LS-1040 |
| Vectashield Vibrance mounting media | Vector labs | H-1700 |

**Kits/Materials**

| | | |
|---|---|---|
| 15 u-Slide VI ibiTreat 6 chamber | Ibidi | 80606 |
| 8 well glass chamber removable slides | Ibidi | 80841 |
| 35 mm round glass bottom | Mattek Corp | P35G-1.5-14C |
| 4-20% Mini Protean TGX stain Free gels | BioRad | 4568094 |
| Clariom D human DNA microarray | Thermo Scientific | 902922 |
| Colorfrost plus microscope slides | Cardinal health | M6148-3P |
| COMET Assay Kit | R&D Systems | 4250-050-K |
| Duolink In Situ Red Starter kit Mouse/Rabbit | Sigma | Duo92101-1KIT |
| Duolink PLA control Kit-PPI | Sigma | Duo92202-1KIT |
| Ion AmpliSeq Transcriptome Mouse Gene Expression Kit | Thermo Scientific | A3655A |
| PVDF membrane | BioRad | 1620177 |
| QuantiTect Reverse Transcription Kit | Qiagen | 205311 |
| Qubit RNA HS Assay Kit | Invitrogen | Q32855 |

| | | |
|---|---|---|
| SimpleChIP Enzymatic Chromatin IP kit( magnetic Beads) | Cell Signaling | 91820 |

**Antibodies**

| | | |
|---|---|---|
| γ-H2AX mouse mAb | Cell Signaling | 80312 |
| γ-H2AX rabbit mAb | Cell Signaling | 9718 |
| ACTA2 Rabbit pAb | Protein Tech | 14395-1-AP |
| Beta Actin Peroxidase Mouse mAb | Sigma | A3854 |
| Cytoglobin Rabbit pAb | Protein Tech | 13317-1-AP |
| Goat anti Mouse Alexaflour 488 | Invitrogen | A-11029 |
| Goat anti Mouse Alexaflour 647 | Invitrogen | A-21236 |
| Goat anti Mouse-HRP | BioRad | 1705047 |
| Goat anti Rabbit Alexaflour 488 | Invitrogen | A-11034 |
| Goat anti Rabbit Alexafluor 594 | Invitrogen | A32740 |
| Goat anti Rabbit-HRP | BioRad | 1705046 |
| Goat Serum | Vector labs | S-1000 |
| HMGB2 mouse mAb | Sigma | WH0003148M5-100UG |
| HMGB2 rabbit pAb | Sigma | HPA053314 |
| HMGB2 rabbit pAb (ChIP) | Sigma | H9789 |
| Mouse IgG isotype control | Vector labs | I-2000 |
| NOX 4 Rabbit pAb | Protein Tech | 14347-1-AP |
| Rabbit IgG isotype Control | Vector labs | I-1000 |
| RTU Animal free blocker and diluent | Vector labs | SP-5035 |

| **Primer List** | **Forward** | **Reverse** |
|---|---|---|
| Human CYGB Ex 2&3 | CAA GGT GGA ACC GGT GTA CT | TCA CGT GGC TGT AGA TGA GG |
| Human NOX 1 (variant 1,2,3) | AAT CCC ATC CAG TCC CGA AA | CCA TGA GAA TCA AGG CTA TT |
| Human NOX 2/CYBB | AGG AAA TAA GGA GAA AAG AG | TGT GTA TAC CTC CTT CAA TT |
| Human NOX 3 | AGG ACA ATA GCA GGC GTG AC | TTT GGC CTC GAA CAA TCC GA |
| Human NOX 4 (variant 4) | CCA CCA GAT GTT GGG GGA TT | CTC CTG GTT CTC CTG CTT GG |
| Human NOX 5 (variant 1) | GCA GAG CGA TTC TTT GCC CT | CTG TCC CGA GTG GGG ATG AA |

### SUPPLEMENTARY TABLE 2

**Merged list of identified cytoglobin interacting protein.**

| Protein names | Gene names | Change | p-value | Majority protein IDs |
|---|---|---|---|---|
| Cytoglobin | CYGB | 13.0 | 0.0000 | K7EMC7;Q8WWM9 |
| Histidine triad nucleotide-binding protein 1 | HINT1 | 4.3 | 0.0005 | D6RD60;D6RE99;D6REP8;P49773 |
| Phosphoglycerate kinase 1 | PGK1 | 4.0 | 0.0030 | P00558;P00558-2 |
| Serine/threonine-protein phosphatase 2A 65 kDa regulatory subunit A alpha isoform | PPP2R1A | 3.9 | 0.0076 | B3KQV6;C9J9C1;P30153 |
| Gamma-soluble NSF attachment protein | NAPG | 3.8 | 0.0000 | Q99747;Q99747-2 |
| Transferrin receptor protein 1 | TFRC | 3.7 | 0.0001 | F8WBE5;P02786 |
| ATP-citrate synthase | ACLY | 3.6 | 0.0001 | P53396;P53396-2;P53396-3 |
| Peptidyl-prolyl cis-trans isomerase FKBP4; Peptidyl-prolyl cis-trans isomerase FKBP4, N-terminally processed | FKBP4 | 3.5 | 0.0045 | Q02790 |
| Histone-lysine N-methyltransferase SETD2 | SETD2 | 3.4 | 0.0029 | A0A1W2PPX9;C9JG86;H7BXT4;H7BZ93;H7C3H4;Q9BYW2;Q9BYW2-2;Q9BYW2-3 |
| Proliferating cell nuclear antigen | PCNA | 3.3 | 0.0040 | P12004 |
| Tubulin beta-3 chain | TUBB3 | 3.2 | 0.0000 | A0A0B4J269;Q13509;Q13509-2 |
| SRSF protein kinase 1 | SRPK1 | 3.1 | 0.0000 | H3BLV9;Q96SB4;Q96SB4-3;Q96SB4-4 |
| Membrane-associated progesterone receptor component 1 | PGRMC1 | 3.1 | 0.0005 | O00264;O00264-2 |
| Probable ATP-dependent RNA helicase DDX10 | DDX10 | 3.1 | 0.0040 | A0A3B3ISR7;E9PIF2;Q13206 |
| Ubiquitin carboxyl-terminal hydrolase | UCHL1 | 3.1 | 0.0041 | D6R956;D6R974;D6RE83;P09936 |
| Glucose-6-phosphate isomerase | GPI | 3.1 | 0.0013 | A0A0A0MTS2;A0A0J9YX90;A0A0J9YXP8;A0A0J9YYH3;A0A2R8Y6C7;A0A2U3TZU2;K7EPY4;K7EQ48;P06744;P06744-2 |
| Phosphoglycerate mutase 1;Probable phosphoglycerate mutase 4 | PGAM1;PGAM4 | 3.0 | 0.0000 | P18669;Q8N0Y7 |
| Protein flightless-1 homolog | FLII | 2.9 | 0.0001 | Q13045-2 |
| | FMR1 | 2.9 | 0.0018 | R9WNI0 |
| Transforming protein RhoA;Rho-related GTP-binding protein RhoC | RHOA;RHOC | 2.9 | 0.0010 | C9JNR4;C9JX21;E9PQH6;P08134;P61586;Q5JR05;Q5JR07;Q5JR08 |
| Leucine--tRNA ligase, cytoplasmic | LARS | 2.8 | 0.0060 | Q9P2J5;Q9P2J5-2;Q9P2J5-3 |
| Trifunctional purine biosynthetic protein adenosine-3 | GART | 2.8 | 0.0011 | C9JBJ1;C9JTV6;F8WD69;P22102;P22102-2 |
| Inorganic pyrophosphatase | PPA1 | 2.6 | 0.0003 | Q15181;Q5SQT6 |
| Adenylosuccinate lyase | ADSL | 2.6 | 0.0013 | A0A096LNY4;A0A096LNY5;A0A096LNY6;A0A096LP72;A0A0A6YY92;A0A1B0GTG9;A0A1B0GTJ7;A0A1B0GW |

| | | | | |
|---|---|---|---|---|
| | | | | F8;A0A1B0GWJ0;A0A1W2PRX2;B4DEP1;P30566;P30566-2 |

**<u>Supplementary Table 2 (continued).</u> Merged list of identified cytoglobin interacting protein.**

| Protein names | Gene names | Change | p-value | Majority protein IDs |
|---|---|---|---|---|
| Vesicle-fusing ATPase | NSF | 2.58199 | 0.002 | I3L0N3;P46459;P46459-2 |
| Guanine nucleotide-binding protein G(I)/G(S)/G(T) subunit beta-1 | GNB1;GNB2 | 2.51214 | 0.004 | B1AKQ8;E7EP32;F6UT28;F6X3N5;P62873;P62873-2 |
| Anaphase-promoting complex subunit 7 | ANAPC7 | 2.49445 | 0.004 | Q9UJX3;Q9UJX3-2 |
| Peroxiredoxin-2 | PRDX2 | 2.40082 | 0.008 | P32119 |
| Cytosolic Fe-S cluster assembly factor NUBP2 | NUBP2 | 2.38043 | 0.002 | B7Z6P0;H3BQR2;Q9Y5Y2 |
| Alpha-enolase | ENO1 | 2.36039 | 0.004 | A0A2R8Y6G6;P06733;P06733-2 |
| Creatine kinase B-type | CKB | 2.12005 | 1E-04 | G3V4N7;P12277 |
| D-3-phosphoglycerate dehydrogenase | PHGDH | 2.11602 | 4E-04 | A0A286YF22;A0A286YF78;A0A286YFA2;A0A286YFL2;A0A286YFM8;A0A2C9F2M7;O43175 |
| Lipoamide acyltransferase component of branched-chain alpha-keto acid dehydrogenase complex | DBT | 2.0609 | 0.004 | P11182;Q5VVL7 |
| Protein cereblon | CRBN | 2.04382 | 2E-06 | A0A1W2PPJ5;J3QT51;J3QT87;Q96SW2;Q96SW2-2 |
| T-complex protein 1 subunit alpha | TCP1 | 2.04164 | 0.008 | E7EQR6;E7ERF2;F5H282;P17987 |
| Synapse-associated protein 1 | SYAP1 | 1.95635 | 0.002 | Q96A49 |
| Actin, cytoplasmic 1 | ACTB | 1.95063 | 0.001 | A0A2R8Y793;P60709 |
| Ras-related proteins | RAB1B;RAB15;RAB8B;RAB1A;RAB10 | 1.92371 | 0.001 | A0A2R8YDI9;A0A2R8YFB8;E7END7;E9PLD0;G3V196;H0YL94;H0YLJ8;H0YMN7;H0YNE9;P51153;P59190;P59190-2;P61006;P61006-2;P61026;P62820;P62820-2;P62820-3;Q92928;Q92930;Q9H0U4 |
| Histone H2A | H2AFV;H2AFZ | 1.92162 | 0.004 | A0A494C189;C9J0D1;P0C0S5;Q71UI9;Q71UI9-2;Q71UI9-3;Q71UI9-4 |
| Sorting nexin-3 | SNX3 | 1.91122 | 4E-06 | O60493;O60493-2;O60493-3;O60493-4 |
| T-complex protein 1 subunit theta | CCT8 | 1.89559 | 0.008 | H7C4C8;P50990;P50990-2;P50990-3 |
| Talin-1 | TLN1 | 1.85292 | 0.003 | Q9Y490 |
| 14-3-3 protein beta/alpha | YWHAB | 1.848 | 0.001 | P31946-2 |
| Nucleosome assembly protein 1-like 1 | NAP1L1 | 1.79909 | 7E-04 | B7Z9C2;F5H4R6;F8VRJ2;F8VUX1;F8VV59;F8VVB5;F8VXI6;F8VY35;F8W020;F8W0J6;F8W118;F8W543;H0YH88;H0YHC3;H0YIV4;P55209;P55209-2;P55209-3 |
| Prohibitin | PHB | 1.7556 | 5E-04 | C9JW96;C9JZ20;E7ESE2;E9PCW0;P35232;P35232-2 |
| Profilin-1 | PFN1 | 1.73129 | 2E-04 | K7EJ44;P07737 |
| Eukaryotic initiation factor 4A-I | EIF4A1;EIF4A2 | 1.71647 | 0.005 | E7EQG2;J3KSZ0;J3KT12;J3KTB5;J3QL43;J3QS69;P60842;P60842-2;Q14240;Q14240-2 |
| Transgelin-2 | TAGLN2 | 1.69279 | 0.005 | P37802;P37802-2;X6RJP6 |
| Guanine nucleotide-binding protein G(s) subunit alpha isoforms | GNAS;GNAO1;GNAI2;GNAT2;GNAL;GNAI3;GNAI1;GNAT3 | 1.69143 | 0.005 | A0A087WTB6;A0A087WZE5;A0A0A0MR13;A0A1W2PQK2;A0A1W2PRE1;A0A1W2PRJ7;A0A1W2PS82;A0A3B3IUA8;A2A2R6;A8MTJ3;H0Y7E8;H0Y7F4;P04899;P04899-4;P08754;P09471;P09471- |

| | | | | |
|---|---|---|---|---|
| | | | | 2;P11488;P19087;P38405;P38405-2;P63092;P63092-2;P63092-3;P63092-4;P63096;Q5JWD1;Q5JWE9;Q5JWF2;Q5JWF2-2 |
| Replication protein A 32 kDa subunit | RPA2 | 1.68494 | 9E-06 | P15927;P15927-2;P15927-3;Q5TEJ7 |
| 39S ribosomal potein L11, mitochondrial | MRPL11 | 1.65812 | 0.003 | Q9Y3B7;Q9Y3B7-2;Q9Y3B7-3 |
| Translation machinery-associated protein 16 | TMA16 | 1.59732 | 0.002 | D6RA57;H0Y9X1;Q96EY4 |
| Clathrin heavy chain;Clathrin heavy chain 1 | CLTC | 1.59677 | 3E-05 | A0A087WVQ6;Q00610;Q00610-2 |
| C-1-tetrahydrofolate synthase, cytoplasmic | MTHFD1 | 1.59265 | 0.001 | A0A384N5Y3;F5H2F4;P11586;V9GYY3;V9GZ78 |
| Actin, cytoplasmic 2;Actin, cytoplasmic 2, N-terminally processed | ACTG1 | 1.54246 | 0.007 | I3L1U9;I3L3I0;P63261 |

### SUPPLEMENTARY TABLE 3

**Table 2. Merged list of identified cytoglobin interacting protein after treatment with hydrogen peroxide**

| Protein names | Gene names | Change | p-value | Majority protein IDs |
|---|---|---|---|---|
| Cytoglobin | CYGB | 14.17 | 0.00000 | K7EMC7;Q8WWM9 |
| Metastasis-associated protein MTA1 | MTA1 | 3.85 | 0.00658 | E7ESY4;H0Y4T7;Q13330 |
| Regulator of nonsense transcripts 2 | UPF2 | 3.54 | 0.00000 | Q9HAU5 |
| 28S ribosomal protein S14, mitochondrial | MRPS14 | 3.50 | 0.00007 | O60783 |
| Gamma-soluble NSF attachment protein | NAPG | 3.32 | 0.00247 | Q99747;Q99747-2 |
| TBC1 domain family member 5 | TBC1D5 | 3.11 | 0.00235 | C9J3F6;Q92609;Q92609-2 |
| High mobility group protein B2 | HMGB2 | 2.99 | 0.00209 | D6R9A6;P26583 |
| 60S ribosomal protein L22-like 1 | RPL22L1 | 2.95 | 0.00004 | C9JYQ9;H0Y8C2;Q6P5R6 |
| Cerebellar degeneration-related protein 2-like | CDR2L | 2.93 | 0.00284 | Q86X02 |
| Protein FRG1 | FRG1 | 2.89 | 0.00103 | E9PLY7;Q14331 |
| 60S ribosomal protein L38 | RPL38 | 2.85 | 0.00038 | J3KT73;J3QL01;P63173 |
| Translation machinery-associated protein 16 | TMA16 | 2.71 | 0.00017 | D6RA57;H0Y9X1;Q96EY4 |
| 39S ribosomal protein L33, mitochondrial | MRPL33 | 2.68 | 0.00065 | O75394 |
| 28S ribosomal protein S12, mitochondrial | MRPS12 | 2.52 | 0.00288 | O15235 |
| Serine/threonine-protein phosphatase 6 regulatory subunit 1 | PPP6R1 | 2.51 | 0.00987 | Q9UPN7 |
| Protein cereblon | CRBN | 2.26 | 0.00098 | A0A1W2PPJ5;J3QT51;J3QT87;Q96SW2;Q96SW2-2 |
| 60S ribosomal protein L22 | RPL22 | 2.16 | 0.00013 | P35268 |
| Actin-related protein 2/3 complex subunit 5 | ARPC5 | 1.92 | 0.00654 | B1ALC0;O15511;O15511-2 |
| Calcium-binding and coiled-coil domain-containing protein 2 | CALCOCO2 | 1.89 | 0.00085 | Q13137;Q13137-2;Q13137-3;Q13137-4;Q13137-5 |
| WD repeat-containing protein 13 | WDR13 | 1.86 | 0.00741 | A0A087X091;Q9H1Z4;Q9H1Z4-2 |
| Non-histone chromosomal protein HMG-17 | HMGN2 | 1.80 | 0.00059 | P05204 |
| Focadhesin | FOCAD | 1.76 | 0.00533 | Q5VW36 |
| E3 ubiquitin-protein ligase HECTD3 | HECTD3 | 1.74 | 0.00560 | Q5T447 |

| | | | | |
|---|---|---|---|---|
| Succinate dehydrogenase [ubiquinone] iron-sulfur subunit, mitochondrial | SDHB | 1.74 | 0.00145 | A0A087WWT1;A0A087WXX8;P21912 |
| Cytosolic non-specific dipeptidase | CNDP2 | 1.64 | 0.00108 | A0A087WYZ1;J3KSV5;J3QKQ0;J3QKT2;J3QLU1;J3QR27;J3QRD0;Q96KP4;Q96KP4-2 |
| PHD finger protein 6 | PHF6 | 1.64 | 0.00048 | A0A0D9SGE8;Q5JRC6;Q8IWS0;Q8IWS0-2;Q8IWS0-3;Q8IWS0-4;Q8IWS0-5 |
| Translation machinery-associated protein 7 | TMA7 | 1.61 | 0.00009 | Q9Y2S6 |
| Tax1-binding protein 1 | TAX1BP1 | 1.61 | 0.00280 | B8ZZD4;Q86VP1;Q86VP1-2;Q86VP1-3;Q86VP1-4 |
| Cell cycle progression protein 1 | CCPG1 | 1.55 | 0.00650 | Q9ULG6;Q9ULG6-2;Q9ULG6-4;Q9ULG6-5 |
| Electron transfer flavoprotein subunit alpha, mitochondrial | ETFA | 1.52 | 0.00604 | H0YK49;H0YKF0;H0YL12;H0YLU7;H0YNX6;P13804;P13804-2 |
| Uncharacterized protein C8orf59 | C8orf59 | 1.51 | 0.00449 | Q8N0T1 |

#### SUPPLEMENTARY FIGURES

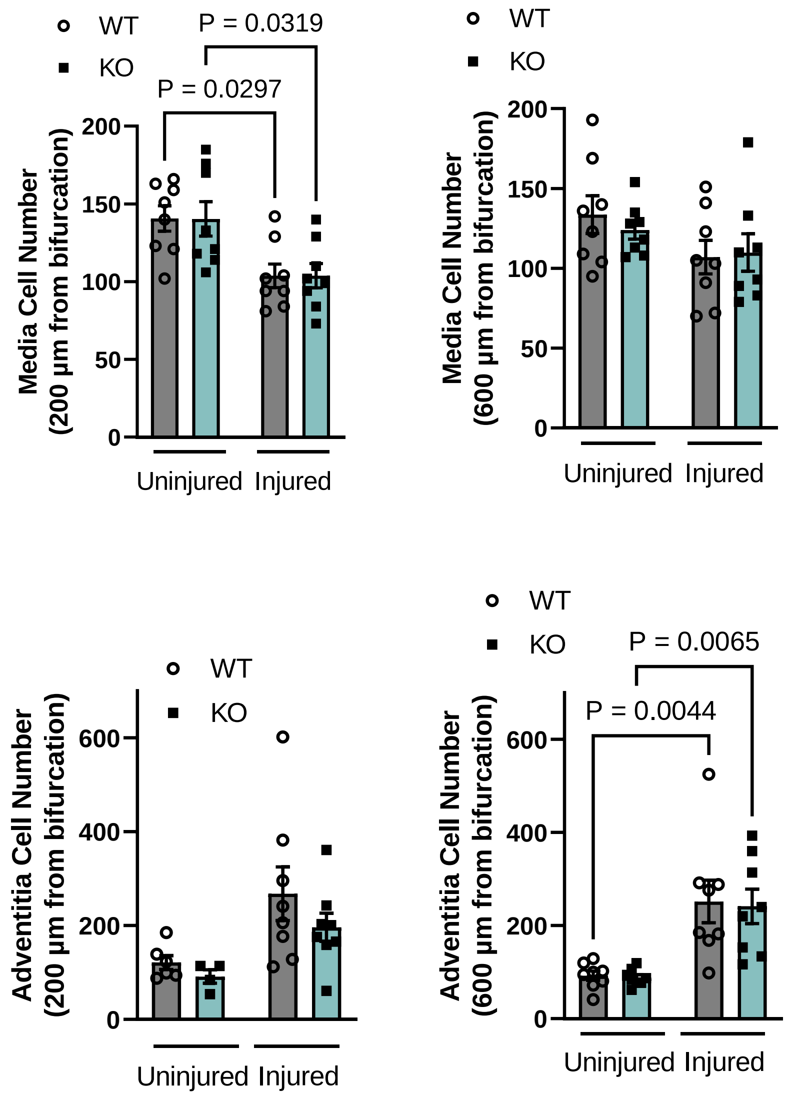


**Supplementary Fig 1. Changes in medial and adventitial cell number in the mouse left common carotid artery, 3-day post-ligation is unaffected by global deletion of cytoglobin.** Medial and adventitial cells were enumerated in tissue sections obtained from the right (uninjured) and left (injured) common carotid arteries, 3-day post-ligation of the left common carotid artery. Both male and females wild-type (WT) and cytoglobin global knockout (KO) mice were used for these experiments. Sections sampled were taken 200 µm from the bifurcation. Each point represents one mice. Results results are presented as +/- SEM and were analyzed by two-way ANOVA, followed by Tukey's post hoc test. Only p values less than 0.05 are shown.

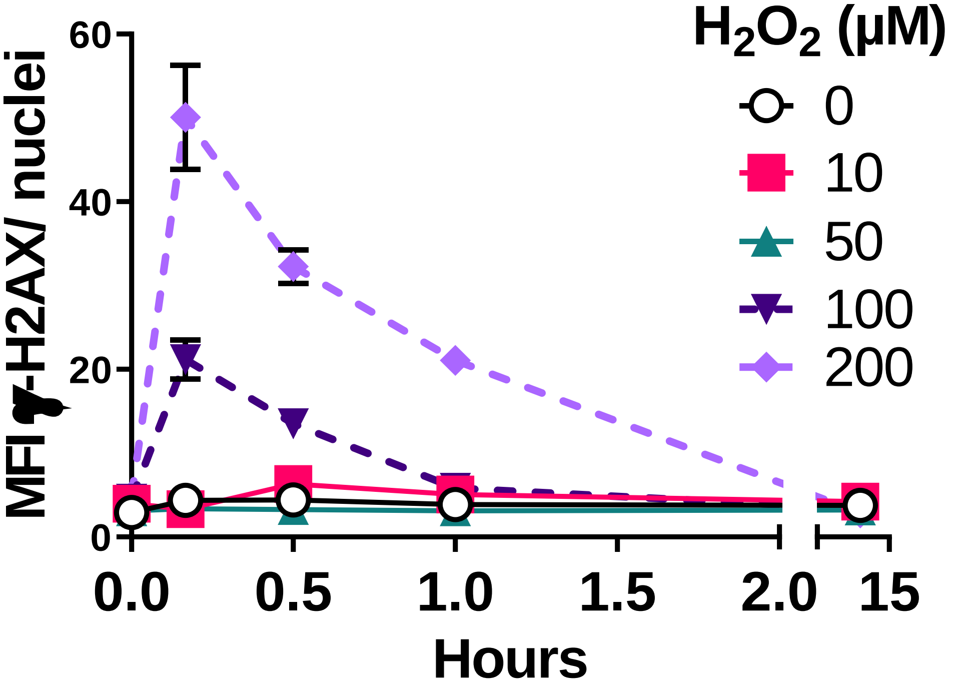


**Supplementary Fig. 2. Hydrogen peroxide transiently increases H2AX phosphorylation (γ-H2AX) in cultured human aortic vascular smooth muscle cells.** Human vascular smooth muscle cells were treated with increasing concentrations of hydrogen peroxide over a 15 hours. Cells were then stained for γ-H2AX and mean fluorescence intensity (MFI) was determined; n=3; results are presented as +/- SEM.

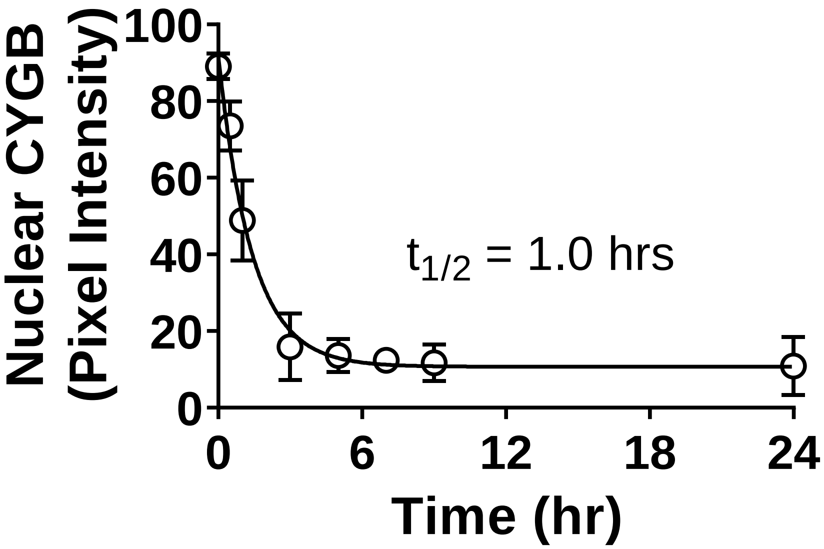


**Supplementary Fig. 3.** Time course of cytoglobin nuclear export over time following incubation of human vascular smooth muscle cells grown in complete media (GM) with 0.2% serum containing serum starvation media (SS). The solid line represents the non-linear regression from a one exponential decay model yielding a half life of 1.0 hours; n = 4. Error bars are SEM.

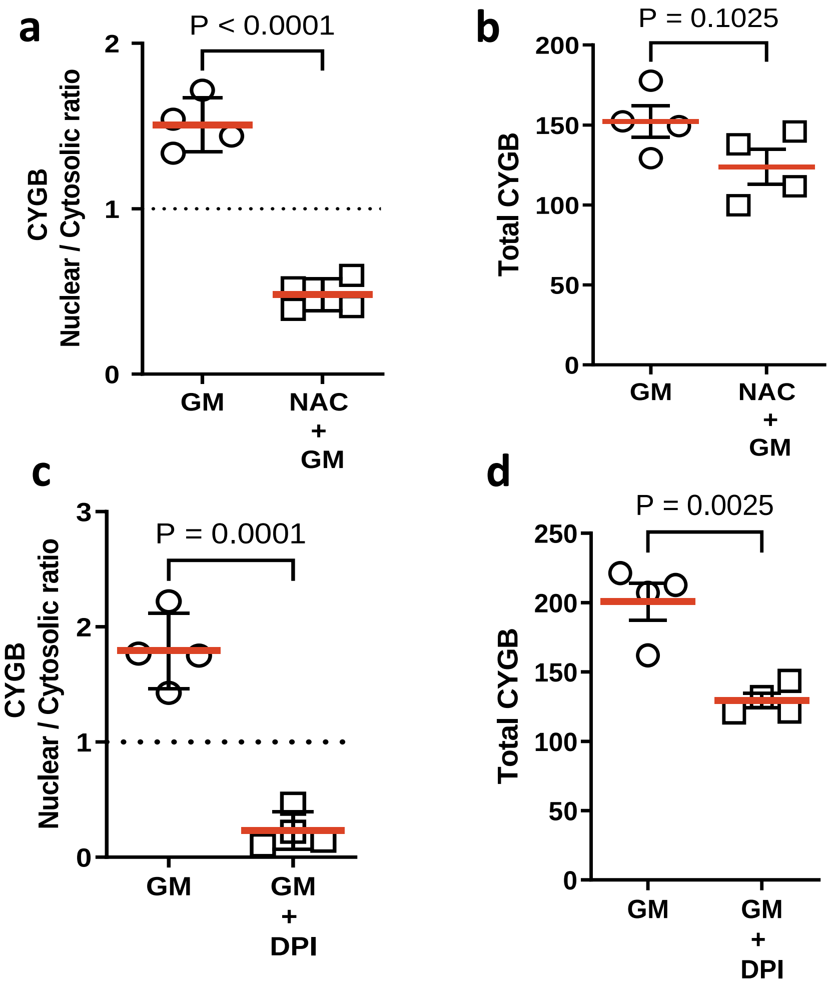


**Supplementary Fig. 4. Antioxidant treatment inhibits the nuclear translocation of cytoglobin. (a)** Human smooth muscle cells were cultured in 0.2% serum for 5 hours and incubated with 1mM N-acetyl cysteine (NAC) in 5% serum for up to 24 hours. Cells were immunostained for CYGB –red and DAPI- blue, visualized by confocal microscopy, and quantitated for the ratio of nuclear to cytosolic CYGB pixel intensity ratio. A ratio greater than 1 (dotted line) is indicative of nuclear localization. Data are presented as +/- SEM. Each dot symbol represents an independent experiment. **(b)** Quantitation of total cytoglobin pixel intensity following NAC treatment. **(c)** Human vascular smooth muscle cells were cultured in 0.2% serum and incubated with 10µM of the NOX inhibitor, diphenyleneiodonium (DPI). **(d)** Quantitative analysis of total CYGB pixel intensity following DPI treatment. Data are presented as mean +/- SEM; p values were determined by Student's t test.

a

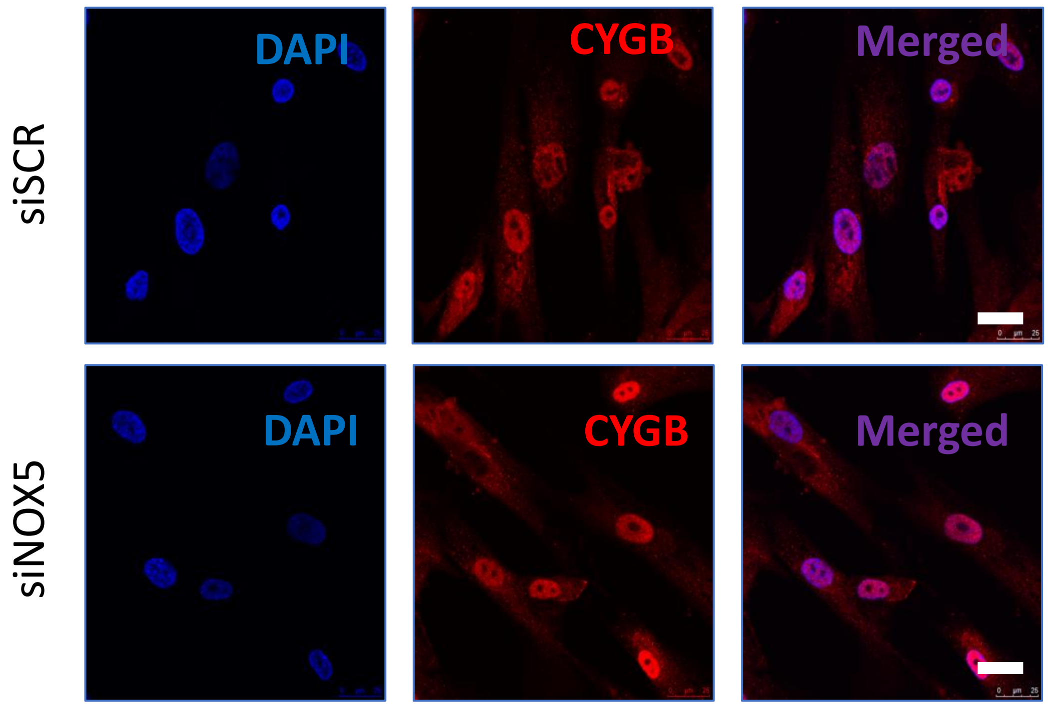


b

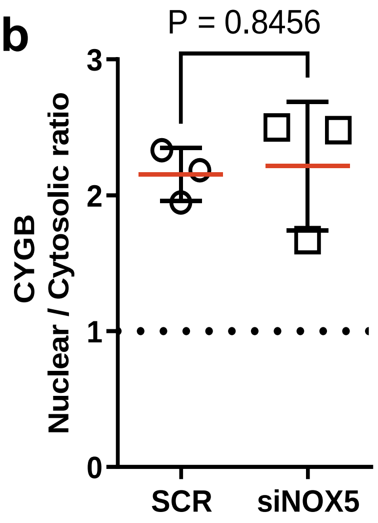


**Supplementary Fig 5. NOX5 silencing does not affect the subcellular localization of cytoglobin in HAoSMCs. (a)** Representative immunofluorescence images of subcultured human vascular smooth muscle cells following silencing of NOX5. CYGB – red and DAPI – blue. **(b)** Quantitative analysis of the nuclear to cytosolic cytoglobin pixel intensity ratio. Data are presented as mean +/- SEM; p value was determined by Student's t test.

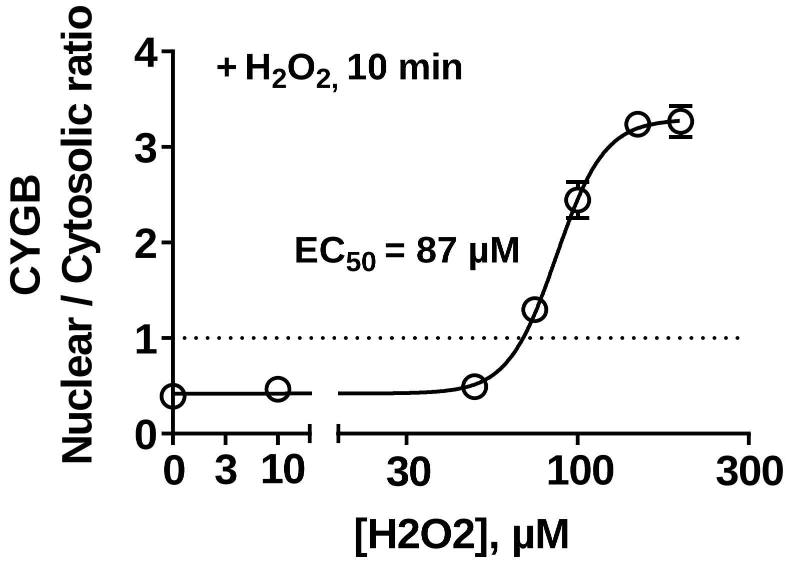


**Supplementary Fig. 6. Hydrogen peroxide stimulates the nuclear accumulation of cytoglobin.** Cultured human vascular smooth muscle cells were treated with increasing concentrations of hydrogen peroxide for 10 min and stained for cytoglobin. The solid line represents the best fit for the non-linear regression of the log-dose response yielding an $EC_{50}$ of 87 µM.; n = 4; results are presented as +/- SEM.

Human coronary vascular smooth muscle cells –
Proximity ligation assay

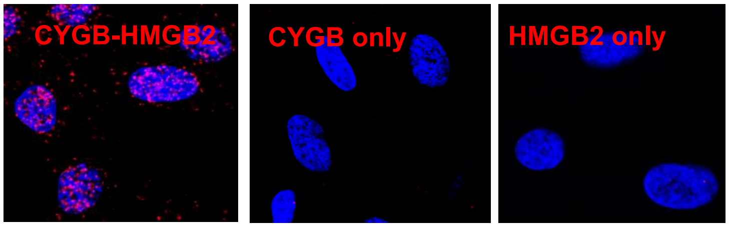


**Supplementary Fig. 7. Cytoglobin and HMGB2 heterodimerizes in human coronary smooth muscle cells *in vitro*.** Proximity ligation assay probing for the interaction of CYGB and HMGB2 in human coronary vascular smooth muscle cells. Bar scale, 25 µm.
